# Spheroid Assembly in Microwells of Defined Geometry for Quantitative Assessment of Aggregation Kinetics and Shape Engineering

**DOI:** 10.1101/2025.09.15.676122

**Authors:** Yuri M. Efremov, Ekaterina Yu. Makarova, Polina I. Koteneva, Daniil O. Golubchikov, Ruslan M. Yanbarisov, Yuri V. Vassilevski, Nastasia V. Kosheleva, Peter S. Timashev

## Abstract

Three-dimensional (3D) cell spheroids are widely used as *in vitro* tissue models, yet quantitative understanding of their morphogenesis remains limited. We present an integrated experimental–computational framework to analyze, model, and modulate the compaction of cell aggregates in agarose microwells of defined geometries. Custom 3D-printed stamps produced circular, square, and triangular microwells of equal cross-sectional area. Time-lapse imaging combined with AI-based segmentation enabled tracking of spheroid morphology, with circularity and projected area serving as quantitative descriptors of compaction. The process followed predictable exponential kinetics, with mesenchymal (HDF) spheroids compacting faster than epithelial (ARPE-19) ones. Computational fluid dynamics (CFD) simulations modeled spheroid rounding as a visco-capillary–driven process, where the extracted visco-capillary velocity unified experimental and simulated dynamics. Mechanical measurements by atomic force microscopy and compression confirmed that differences in surface tension predominantly governed the observed kinetics. Pharmacological modulation of cytoskeletal tension revealed that inhibition of contractility markedly altered spheroid formation dynamics, enabling the generation of stable, non-spherical aggregates. Using this principle as a shape-engineering strategy, we produced aggregates with distinct geometries (brick-like, prismatic, and star-shaped), characterized by an increased surface-to-volume ratio compared to conventional spheroids. Limitations of the approach include the use of pharmacological cytoskeletal modulation and constraints in geometric fidelity arising from printing resolution, agarose casting, cell filling, and intrinsic smoothing of sharp features during cell aggregation. Collectively, this work establishes a geometry-controlled platform for quantitative analysis of spheroid formation and mechanical behavior, and provides a versatile framework for designing cell aggregates with defined shapes.

## Introduction

Three-dimensional (3D) cell spheroids have become widely adopted as *in vitro* models in biomedical research due to their ability to more accurately replicate the physiological characteristics of tissues compared to traditional two-dimensional (2D) cell cultures [1–4]. Unlike monolayer cultures, 3D spheroids foster more natural cell–cell and cell–matrix interactions, leading to gradients of nutrients, oxygen, and signaling molecules that closely mimic the *in vivo* microenvironment. This makes them particularly valuable in tissue engineering, where they serve as building blocks for constructing larger tissue structures [5–7]; in cancer research, where they model tumor growth, invasion, and drug resistance [8,9]; and in drug screening, where they better represent responses to therapeutic compounds [10–12].

Understanding the dynamics of spheroid formation is essential for advancing both fundamental biology and applied biomedical research. The self-assembly of cells into spheroids involves a complex interplay of cell–cell adhesion, migration, proliferation, and extracellular matrix (ECM) remodeling — processes that collectively determine the structural integrity and function of the resulting 3D construct [13–15]. Mechanical properties such as stiffness, viscoelasticity, and surface tension not only indicate spheroid maturation but also actively regulate key cellular behaviors, including differentiation, morphogenesis, nutrient and drug diffusion, and responses to external stimuli [16–18]. These biophysical characteristics can also recapitulate aspects of tumor progression and tissue development. Therefore, gaining a comprehensive understanding of how spheroids form and how their mechanical features impact this process is crucial for optimizing tissue engineering strategies, improving bioprinting fidelity, enhancing drug screening platforms, and developing predictive *in vitro* systems that more accurately model the *in vivo* environment.

Among mechanical parameters, the interplay between surface tension and the effective viscosity of cell aggregates has been shown to determine spheroid formation, maturation, and fusion [17,19–21]. Steinberg et al. introduced a conceptual analogy between liquids and tissues, demonstrating that tissues exhibit measurable surface tension and viscosity [22–25]. The visco-capillary velocity, defined as the ratio of surface tension to viscosity, was proposed as a key governing parameter in many biological processes [21,26–28]. Knowing this parameter is therefore essential for insights into embryonic development, tissue remodeling, cancer invasion, and organ bioprinting. Several techniques have been developed to measure these mechanical properties, either independently or in combination, including compression assays, micropipette aspiration, and others [17,24,29,30]. However, these methods often require specialized equipment and are not easily scalable. There remains a need for more robust, quantitative, and high-throughput techniques. Additionally, the relationship between mechanical parameters measured in fully formed spheroids and those governing the dynamic process of spheroid assembly is still not fully understood.

Modeling approaches have been developed to capture the fundamental dynamics of spheroid behavior, including formation and fusion [26,31,32]. Recent advances in the simulation of viscoelastic fluids have opened new avenues in this field. Unlike purely Newtonian fluid models, viscoelastic formulations account for both the fluid-like and solid-like behaviors characteristic of spheroids, which are composed of densely packed, actively contracting cells. Although these models operate at the continuum level and are relatively simplified, they offer valuable insights into the mechanical regulation of tissue morphogenesis, tumor development, and the engineering of biomimetic tissue constructs. As such, they serve as a promising framework for integrating mechanical principles into the design and interpretation of 3D multicellular systems.

The aim of the current study was to develop a method for extracting the visco-capillary velocity (*ν_p_*) from dynamic observations of spheroid formation and to evaluate the applicability of viscous fluid models incorporating surface tension in describing this process. To accomplish this, non-adhesive agarose microwells with non-circular cross-sections (square and triangular) were fabricated, enabling precise tracking of the dynamic rounding behavior of cell aggregates. Spheroids composed of two distinct cell types, epithelial and mesenchymal, were analyzed using AI-based image segmentation, revealing differences in the kinetics of formation between the two. Additionally, the effects of several pharmacological inhibitors, potentially modulating the visco-capillary velocity, were investigated. A computational fluid dynamics (CFD) model, representing the spheroid as a Newtonian fluid with surface tension, was employed to simulate the rounding process within the shaped microwells. The model showed satisfactory agreement with experimental observations, supporting the relevance of continuum fluid descriptions in capturing key aspects of spheroid morphogenesis, and providing a means of measuring the visco-capillary velocity in the conducted experiments. Furthermore, we demonstrate that pharmacological modulation of this parameter provides a strategy to generate spheroids with predefined, microwell geometry-dependent shapes, thereby establishing a controllable platform for engineering multicellular aggregates with tailored morphological properties. Using this approach, we constructed several types of shape-engineered spheroids (brick-like, prismatic, and star-shaped) and quantitatively characterized their geometric features. In addition, the reversibility of the induced shape changes was evaluated.

## Materials and Methods

### Design of differently-shaped microwells

The stamps were designed using FreeCAD (version 1.0.0) and exported as .stl files (Supplementary Model Files 1–3). Each stamp consisted of a tubular post base, dimensioned to fit securely into the wells of a standard 24-well culture plate. The post was attached to a stopper plate, which allowed precise placement and maintained the stamp at a fixed height above the bottom of the well (Fig. 1A–E). To facilitate air displacement and prevent bubble formation during agarose addition, side cuts were incorporated into the stamp design. At the end of the tubular post, three types of protrusions (approximately 160 ± 3 per stamp) were arranged in a regular square grid to form microwells in the agarose: cylinders (circular cross-section), rectangular cuboids (square cross-section), and equilateral triangular prisms (triangular cross-section). Each protrusion had a height of 0.75 mm, and their lateral dimensions were adjusted to maintain a consistent cross- sectional area of 0.16 mm^2^ (Fig. 1). In the following text, the stamps are referred to as “circular,” “square”, and “triangular” based on the resulting agarose microwell shapes.

**Figure 1.**
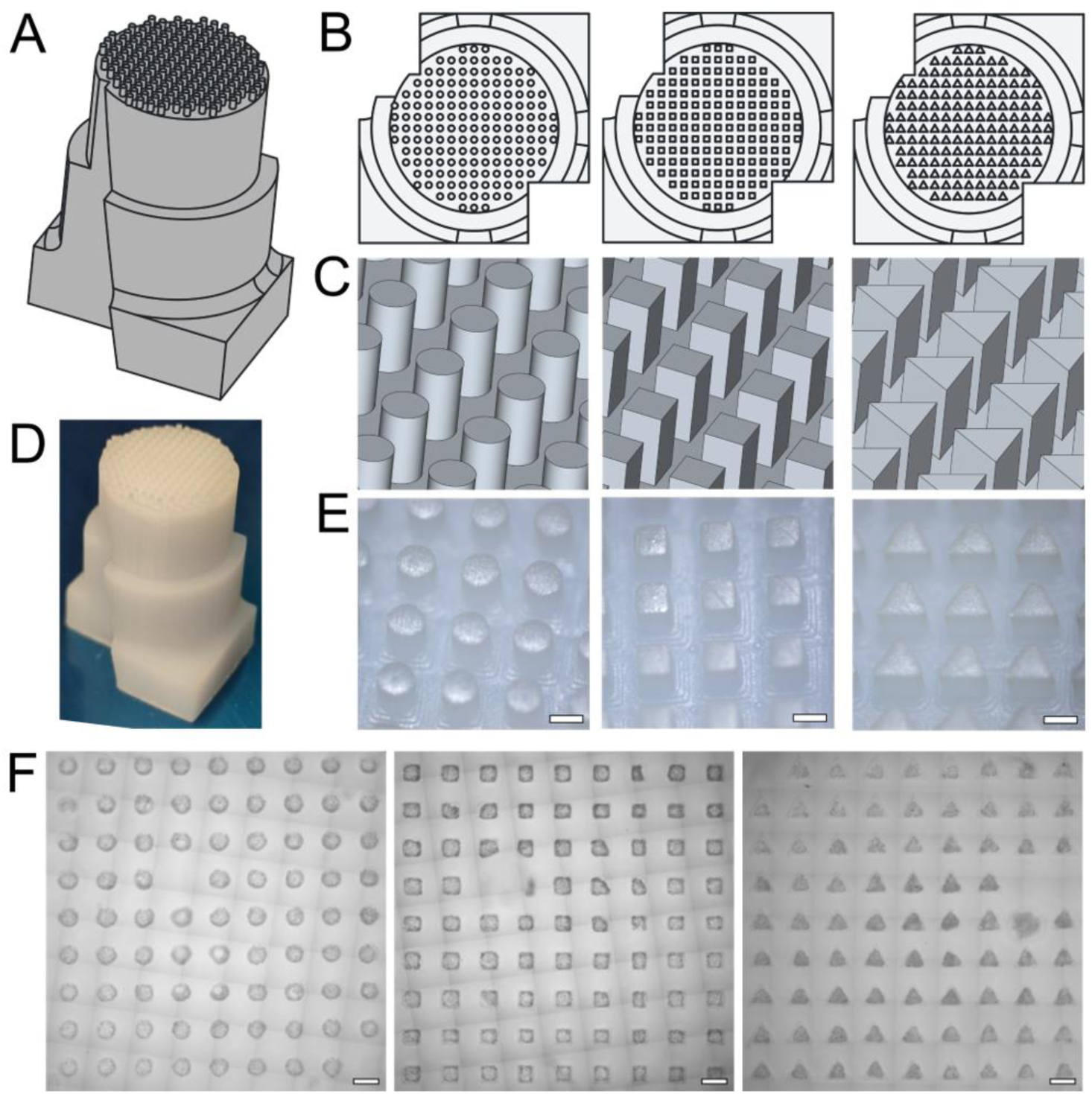
Design of stamps with circular, square, and triangular protrusions. (A) General view of the CAD-created model; (B) Top view of the CAD-created models with different types of protrusions (circular, square, and triangular); (C) Magnified view of the protrusions; (D) Example of a printed stamp; (E) Top part of printed stamps under a stereomicroscope; (F) Example of the microwells formed in agarose in a well of a 24-well plate right after cell seeding. Part of the stitched image is presented. Scale bars are 500 μm.

The stamp models were sliced using Chitubox Basic (version 2.3.1) and printed on an Anycubic Photon D2 DLP printer (Anycubic, China) using Anycubic Tough Resin at a resolution of 51 μm (pixel size). Following printing, the stamps were rinsed in isopropanol for 30 minutes, dried at room temperature for 20 minutes, and UV-cured for an additional 20 minutes. Prior to use, the stamps were sterilized by UV irradiation in a Microcid sterilization chamber (LLC Electronic Medicine, Moscow) for 20 minutes.

### Preparation of microwells

A 3% (w/v) low melting point agarose solution (Invitrogen, Carlsbad, CA) was prepared in Dulbecco’s Modified Eagle Medium (DMEM). The solution was thoroughly mixed and heated in a microwave until the agarose was fully dissolved and liquefied (80–90 °C). Agarose microwells were fabricated under sterile conditions by pipetting 350 μL of the melted agarose into each well of a 24-well tissue culture-treated plate (Falcon, Corning, NY), followed by immediate insertion of the prepared stamps. The agarose was allowed to solidify on the cooling surface (10 °C) of a Dry Bath DB100C (JOANLAB, China) for 5 minutes, after which the stamps were manually removed. To maintain hydration of the microwells until cell seeding, 0.5 mL of DMEM was added to each well.

### Cells and cell spheroid preparation

Two cell lines were used: immortalized human dermal fibroblasts (hTERT-HDFa) and the epithelial cell line ARPE-19, both obtained from the Cell Culture Collection of the N.K. Koltzov Institute of Developmental Biology, Russian Academy of Sciences (RAS). Cells were cultured under standard conditions (37 °C, 95% humidity, 5% CO_2_) in DMEM/F12 medium supplemented with 10% fetal bovine serum (FBS; Thermo Fisher Scientific, USA) and 1% antibiotic–antimycotic solution (Thermo Fisher Scientific, USA).

Upon reaching confluency, cells were detached using Trypsin–EDTA (0.25%, Servicebio, China) and resuspended in fresh medium at the appropriate concentration. To achieve a ratio of 2,000 cells per spheroid, 500 μL of cell suspension containing 3.2 × 10^5^ cells was added to each well of the plate containing the agarose microwells. The plates were centrifuged at 100 × g for 5 minutes and then transferred either to a cell incubator or to the CellInsight CX7 High-Content Screening (HCS) Platform (Thermo Fisher Scientific) for time-lapse imaging of spheroid formation.

To assess cell viability in spheroids, a Live/Dead staining solution was prepared in cell culture medium containing calcein-AM (1 µM, Sigma-Aldrich, Germany), propidium iodide (PI, 2 µM), and Hoechst 33342 (5 µM, Thermo Scientific, USA). The culture medium in agarose microwells was replaced with an equal volume of the staining solution, and samples were incubated for 1 h in an incubator. Subsequently, spheroids were gently extracted from the agarose wells and transferred to glass-bottom dishes (WillCo Wells B.V., Amsterdam, Netherlands) containing FluoroBrite DMEM medium (Thermo Scientific, USA). Imaging was performed using an FV3000 laser scanning confocal microscope (Olympus, Japan) equipped with 405, 488, and 594 nm excitation lasers. Image analysis was conducted in ImageJ: total cell counts were determined from Hoechst 33342-stained nuclei, dead cells were identified by PI staining, and viability was confirmed by calcein-AM fluorescence.

When required, spheroids were gently extracted from the microwells by flushing and transferred into wells of a 6-well tissue culture-treated plate to allow adhesion and cell outgrowth. Phase-contrast images of cell spreading were acquired using an Axio Vert A1 inverted microscope (Carl Zeiss, Germany). The area of cell outgrowth was manually delineated using the Freehand Selection tool in ImageJ (NIH, USA), and the enclosed area was quantified.

### Time-lapse imaging and image processing

Time-lapse imaging was performed using the CellInsight CX7 High-Content Screening (HCS) Platform (Thermo Fisher Scientific) under controlled conditions (37 °C, 80% humidity, and 5% CO₂). Brightfield images were acquired using a 10 × objective across the full area of selected wells, with a time interval of 4 hours between frames ([Movie S1](https://drive.google.com/file/d/1hn8QC8-5QQMhC0lNoePuxm8VPHBmftED/view?usp=drive_link)). In some experiments, a shorter time step (1–1.5 hours) was used to capture the dynamics of fast-forming HDF spheroids.

In experiments with pharmacological treatments, the acting agents (Cytochalasin D (cytD), 1 mM in DMSO; (-)-Blebbistatin (10 mM in DMSO, Sigma, USA); Y-27632 (10 mM in DMSO, Sigma, USA)) were added to the wells with cells seeded in the microwells. Working concentrations were 0.5 μM (for ARPE-19 cells) and 1 μM (for HDF cells) for cytD; 10 μM for Blebbistatin; and 50 μM for Y-27632. Control wells received the maximum equivalent concentration of DMSO (0.5%) corresponding to the highest solvent concentration used in the treatment groups.

Subsequent image processing was conducted in ImageJ (NIH, Bethesda, USA). First, image tiles within each well were stitched into a single composite frame using the Grid/Collection Stitching plugin [33]. Spheroid segmentation was then performed using the SAMJ-IJ plugin (Segment-anything-models-java for ImageJ plugin, GitHub, https://github.com/segment-anything-models-java/SAMJ-IJ), which applies SAM-2 (Segment Anything Model 2) [34]. This process was repeated for each time frame in the time-lapse series with a custom-written ImageJ script. For the initial frame, points were manually selected within the cell-filled microwells to serve as prompts (seed locations) for segmentation. These prompted the plugin to detect and delineate the spheroid boundary, producing regions of interest (ROIs). The centroid (center of mass) of each segmented ROI was calculated using ImageJ’s built-in measurement tools and used as the updated prompt for the next frame, allowing the method to track changes in spheroid position and morphology over time. ROIs were stored for each time point, enabling the time-dependent measurement of morphometric parameters ([Movie S3](https://drive.google.com/file/d/1vBilb9kSTcGeOiB63aUwsseH9LZMbGdK/view)). The code used for analysis is publicly available on GitHub: https://github.com/yu-efremov/cell-aggregation-kinetics-openfoam.

ROIs corresponding to spheroid contours were resampled uniformly along arc length to a fixed number of points (1000 points per contour) to ensure consistent spatial resolution across spheroids and time points. Contours corresponding to partially filled wells or segmentation artifacts were excluded using a robust outlier detection procedure based on the median absolute deviation (MAD) of projected area and perimeter. For each parameter, the median value across contours was computed, and deviations from the median were scaled by the corresponding MAD. Contours were rejected if either projected area or perimeter deviated from the group median by more than three scaled MAD units. Visual assessment confirmed that excluded spheroids corresponded to partially filled microwells (low projected area), morphologically abnormal spheroids (low or high perimeter), or segmentation errors (high perimeter).

The following morphological parameters were extracted for each spheroid contour: projected area, circularity, roundness, and solidity. The formula for circularity is 4π(area/perimeter^2^), where a value of 1.0 represents a perfect circle. Roundness is calculated as 4*area/(π*major_axis^2^), and solidity is the ratio of area to convex hull area of the ROI. Perimeter-to-normalized area was calculated by first scaling each contour to remove size differences, using the equivalent radius derived from its projected area. Circularity and projected area time dependencies were fitted with exponential rise-to-maximum and exponential decay functions, respectively:

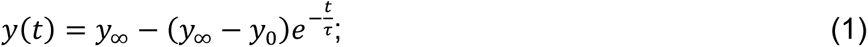

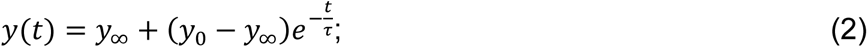

where *y(t)* is the measured parameter, *y_0_* is the initial value, *y_∞_* is the plateau value, and τ is the characteristic time constant. For circularity, the initial time points showing a small transient decrease were excluded from the fit, as described in the Results.

For endpoint analyses of treated and shape-engineered spheroids, contours were centered and rotationally aligned by maximizing the circular cross-correlation of their radial distance functions to ensure consistent angular orientation prior to averaging and curvature analysis. Contours were smoothed using a Savitzky–Golay filter (window length = 100 points, polynomial order = 3) to reduce high-frequency noise while preserving corner features. Local boundary curvature (κ) was then computed using discrete differential geometry, with first- and second-order spatial derivatives estimated via central finite differences. From the curvature profile, curvature variance (Var(κ)) and a characteristic maximum curvature (κ_max_) were extracted. Curvature variance was used to quantify deviation from circular geometry, as a perfect circle exhibits constant curvature. To obtain a noise-robust estimate of corner sharpness, κ_max_ was defined as the mean of the top 5% of the highest absolute curvature values.

### Computational fluid dynamics simulations

OpenFOAM (v12, OpenFOAM Foundation), an open-source computational fluid dynamics platform, was used for the numerical simulations of the spheroid formation process. It was assumed that the behavior of cells during spheroid formation in the surrounding medium can be described as the flow of two incompressible and immiscible Newtonian fluids (or phases), with the given surface tension coefficient between them and with different viscosities. The Volume of fluid (VoF) method for phase-fraction based interface capturing, which is implemented in the InterFoam, OpenFOAM solver, was used. A uniform 100×100×100 3D cubic mesh (based on a prior mesh sensitivity study, Fig. S1) with the domain size of 1200×1200×1200 μm was used in all simulations with the cell phase occupying the central area. No-slip wall boundary conditions (zero velocity) were prescribed on all domain boundaries. In all simulations, we set the same fixed density for both phases, ⍴ = 1000 kg·m^-3^, and fixed viscosities for the surrounding medium (η_medium_ = 10^-3^ Pa·s) and spheroid cells (η_cells_ = 10^7^ Pa·s), and varied the surface tension coefficient to obtain different visco-capillary velocities. The total simulation time was set to match the duration of the corresponding experiment (120 h). The time step was selected adaptively, constrained by a Courant number condition (Co ≤ 1) and a maximum allowable time step of 0.02 h, as determined from a prior time-step sensitivity analysis. The OpenFOAM case files are publicly available on GitHub: https://github.com/yu-efremov/cell-aggregation-kinetics-openfoam.

### Surface tension estimation by atomic force microscopy and parallel plate compression

The surface tension was estimated based on an approach established in our previous work [35]. Briefly, atomic force microscopy (AFM) provides information on the local elasticity of the outer layer, while compression reflects the global mechanical response of a spheroid. Surface tension was then isolated by analyzing how much of the total compressive resistance arises from surface forces rather than bulk deformation. The detailed procedure is presented in the Supplementary Information.

AFM measurements were performed on live spheroids in growth medium using a Bioscope Resolve system (Bruker, Santa Barbara, CA) mounted on an Axio Observer inverted optical microscope (Carl Zeiss, Germany). The sample temperature was maintained at 37 °C throughout the experiments using the heated stage. Spheroids were transferred in growth medium onto cell culture Petri dishes and were incubated for 1-2 hours to promote adhesion prior to measurements. Due to the spheroidal geometry, AFM measurements were taken from the central top region of each spheroid, where the surface was relatively flat. For nanomechanical mapping, PeakForce QNM-Live Cell probes (PFQNM-LC-A-CAL, Bruker AFM Probes, Camarillo, CA, USA) were used. These probes feature short paddle-shaped cantilevers with pre-calibrated spring constants in the range of 0.09–0.11 N/m and 17-μm-long tips with a nominal radius of 70 nm. Fast Force Volume mode was employed to acquire nanomechanical maps, with spatial resolution of 40x40 points over area of 60 × 60 μm^2^. Force curves were acquired at a vertical piezo speed of 180 μm/s (30 Hz ramp rate, 3 μm ramp size) with a force setpoint of 0.5–1 nN, resulting in indentation depths of around 500 nm. The individual force curves of the force volume maps were processed with the Python scripts, as described previously, using the Hertz’s model to obtain the effective Young’s modulus and standard linear solid model to obtain the relaxation time [36].

Parallel-plate compression was performed using a micro-scale testing system (MicroTester G2, CellScale Biomaterials Testing, Canada). Spheroids were compressed between a fixed, rigid substrate and a displaceable cantilever beam with an attached flat platform in a PBS-filled bath. Compression up to 30% of the initial spheroid height with a speed of 10 µm/s was performed while simultaneously recording the force, upper plate position, and the spheroid shape by a side-view camera. At least 10 spheroids per cell type were measured. The mechanical model for spheroid compression from [35] was then applied to estimate the bulk elastic modulus and the relaxation time. Then, by comparing the surface and bulk elasticity, the effective surface tension was estimated based on the model suggested in the work [37]. The detailed description of the applied models is presented in the Supporting Information.

### Confocal microscopy

The spheroids were fixed in 4% PFA overnight at +4 °C. After three washes in PBS, samples were permeabilized with 0.1% Triton X-100 (Sigma-Aldrich, Saint Louis, Missouri, USA) for 30 min at room temperature. Following three additional PBS washes, non-specific binding was blocked with 10% FBS (Gibco, Thermo Fisher Scientific, Waltham, MA, USA) for 30 min at room temperature. Samples were divided into two groups. The first group was stained with rhodamine–phalloidin (Thermo Fisher Scientific, Waltham, MA, USA) to visualize F-actin (overnight incubation), washed, and mounted in FUnGI optical clearing medium [38]. The second group (ARPE-19 spheroids only) was incubated with primary antibodies against ZO-1 (polyclonal antibody, 1:125, Invitrogen, Carlsbad, CA, USA) diluted in PBS supplemented with 10% FBS and 1% Tween-20 (neoFroxx, Germany) for 24 h at +4 °C. After three PBS washes, spheroids were incubated with Alexa Fluor 488–conjugated secondary antibodies (1:500, Invitrogen, Carlsbad, CA, USA) for 2 h at +4 °C. Then samples were washed and mounted using ProLong Diamond Antifade Mountant with DAPI (Thermo Fisher Scientific, Waltham, MA, USA). Samples from both groups were imaged with an Olympus IX83 laser scanning confocal microscope (Olympus, Japan) equipped with 405 nm, 488 nm, and 594 nm excitation lasers. Z-stacks spanning the entire spheroid thickness were acquired for the first group to quantify spheroid height using ImageJ (NIH, Bethesda, MD, USA).

### Statistical analysis

All experiments were performed with a minimum of two independent biological repeats. The total number of spheroids analyzed after applying exclusion criteria is indicated as *n* for each condition. Statistical significance for measured parameters was evaluated using the Kruskal–Wallis test followed by Dunn’s multiple comparisons test with Bonferroni correction, and by the Mann–Whitney U test for two-group comparisons. Significance levels are indicated in the figures as * for p < 0.05, ** for p < 0.01, and *** for p < 0.001. To estimate uncertainty in characteristic fit times, a nonparametric bootstrap procedure was applied. For each treatment group, individual spheroid time series of projected area or circularity were resampled with replacement and averaged at each time point. The averaged trajectories were then fitted with an exponential model to extract the characteristic time constant. This procedure was repeated 2,000 times to generate bootstrap distributions of the fitted time constants and to estimate their variability.

The uncertainty of derived parameters (e.g., effective viscosity) was calculated using nonparametric bootstrap resampling (2,000 iterations), propagating variability from independently measured quantities. Bootstrap results are presented as median ± half-width of the 95% confidence interval (CI). Between-group comparisons of fitted time constants were performed using a nonparametric permutation test (20,000 label permutations). Under the null hypothesis of exchangeability between groups, treatment labels were randomly permuted and the difference in the median τ was recalculated to generate a null distribution. The achieved significance level (ASL) was computed as the proportion of permuted differences whose absolute value was greater than or equal to the observed difference.

## Results

### Spheroid formation in wells with different shapes

Microwells with three distinct cross-sectional geometries (circular, square, and equilateral triangular) were fabricated using a combination of 3D printing and agarose stamping [39–41]. All microwell types were designed to have the same cross-sectional area (0.16 mm^2^) and a uniform depth of 0.75 mm (Figs. 1 and 2). Mesenchymal (HDF) and epithelial (ARPE-19) cells were seeded into these microwells, and the spheroid formation process was monitored using time-lapse imaging at regular 4-hour intervals.

**Figure 2.**
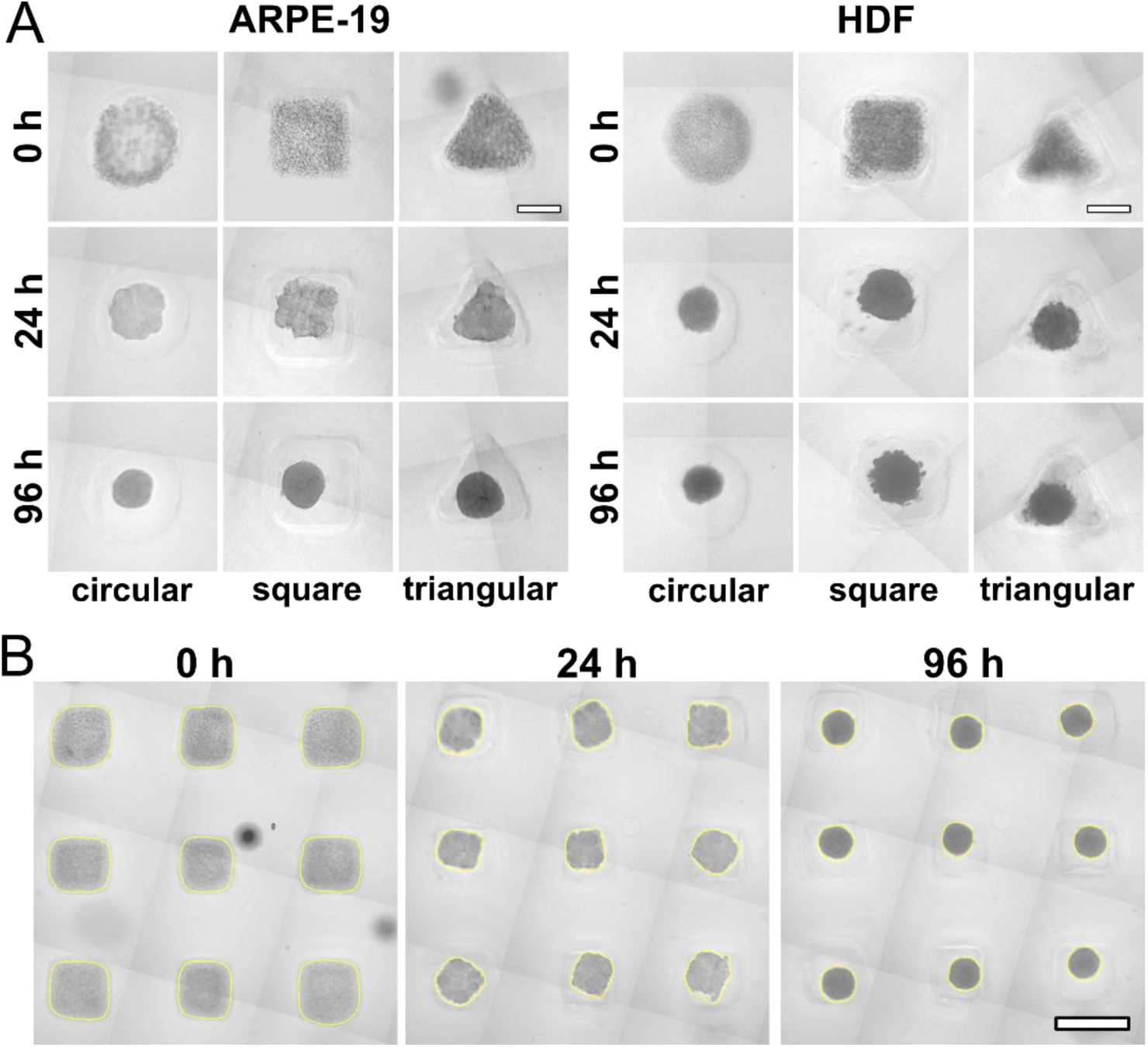
Time-resolved observation of spheroid formation from ARPE-19 and HDF cells in microwells of different cross-sectional geometries (circular, square, triangular). **(A)** The same microwells are shown at three time points (0 h, 24 h, and 96 h) from a continuous series with a 4 h interval. Scale bars are 200 μm. (**B**) Example of SAM-based segmentation of ARPE-19 cell aggregates in square microwells (3×3 microwell subset, yellow outlines). Shown are the time points at 0 h, 24 h, and 96 h. Scale bars are 500 μm.

This setup enabled the simultaneous acquisition of morphological data from a large number of spheroids, approximately 100 spheroids per well across multiple wells in a single plate, making the approach suitable for high-throughput observation. Remarkably, in all three microwell geometries, cell aggregates underwent compaction and ultimately adopted a spherical shape, as indicated by circular 2D projections in brightfield images (Fig. 2A, [Movie S1](https://drive.google.com/file/d/1hn8QC8-5QQMhC0lNoePuxm8VPHBmftED/view), [Movie S2](https://drive.google.com/file/d/1ZLSKOM2nSiighMnBMKWZEG5pS4aC0Ojs/view)). However, the kinetics of compaction differed between cell types: HDF spheroids compacted and reached a spherical morphology drastically faster than ARPE-19 spheroids (at ∼10–20 h vs ∼60–80 h), suggesting cell-type-specific differences in the observed process and visco-capillary behavior.

To quantify differences in the spheroid formation process, image analysis was performed on the time-lapse dataset. Spheroid outlines were identified using the Segment Anything Model (SAM), implemented via the SAMJ-IJ plugin in Fiji/ImageJ. The analysis was designed to track the morphological evolution of each individual spheroid over time. Examples of segmented spheroids with identified outlines are shown in Fig. 2B and [Movie S3](https://drive.google.com/file/d/1vBilb9kSTcGeOiB63aUwsseH9LZMbGdK/view). SAMJ-IJ–based segmentation was validated against manual segmentation on a representative dataset comprising 20 spheroids across five time points, demonstrating sufficient agreement between the two methods (Fig. S2 and Supplementary Text). Agreement was assessed both visually and quantitatively using standard metrics, including the Dice coefficient and Intersection-over-Union (IoU) for mask overlap, as well as Bland–Altman analysis for projected area and circularity. Additionally, the segmentation results were robust to variations in the initial prompt location: small changes in the user-defined seed point did not noticeably affect the extracted contours or derived morphometric parameters (Fig. S2). Partially filled microwells and morphologically abnormal spheroids were excluded from analysis using threshold criteria based on projected surface area and perimeter, as detailed in the Methods section.

To assess the effective microwell geometry resulting from the combined effects of stamp fabrication, agarose casting, and cell seeding, we performed shape analysis of the initial cell-filled microwell contours and compared them with the nominal CAD geometries using Dice similarity coefficients and Hausdorff distances (Fig. S3 and Supplementary Text). Dice scores were high for square and circular microwells (0.91 ± 0.03, n = 137, and 0.94 ± 0.02, n = 148, respectively), indicating strong global shape correspondence, but lower for triangular microwells (0.76 ± 0.03, n = 130). Deviations were primarily localized at sharp corners, particularly in triangular microwells, where rounding consistent with printing and casting limits was observed. These local discrepancies were reflected in the Hausdorff distance values, which were low for circular (38 ± 8 µm) and square (45 ± 10 µm) geometries and higher for triangular geometries (118 ± 11 µm), as expected due to their sharper features.

From the segmented outlines, the following morphological parameters were computed and plotted as a function of time: projected area, circularity, roundness, and solidity (Fig. 3). In all microwell types, the projected area decreased over time, with faster compaction observed for mesenchymal HDF spheroids. In non-circular microwells (square and triangular), shape descriptors such as circularity, roundness, and solidity increased over time, indicating a transition toward a more spherical morphology. Among these, circularity exhibited the most pronounced change. In circular microwells, however, only weak changes in circularity were observed, as the aggregates already adopted a circular projection from the outset.

**Figure 3.**
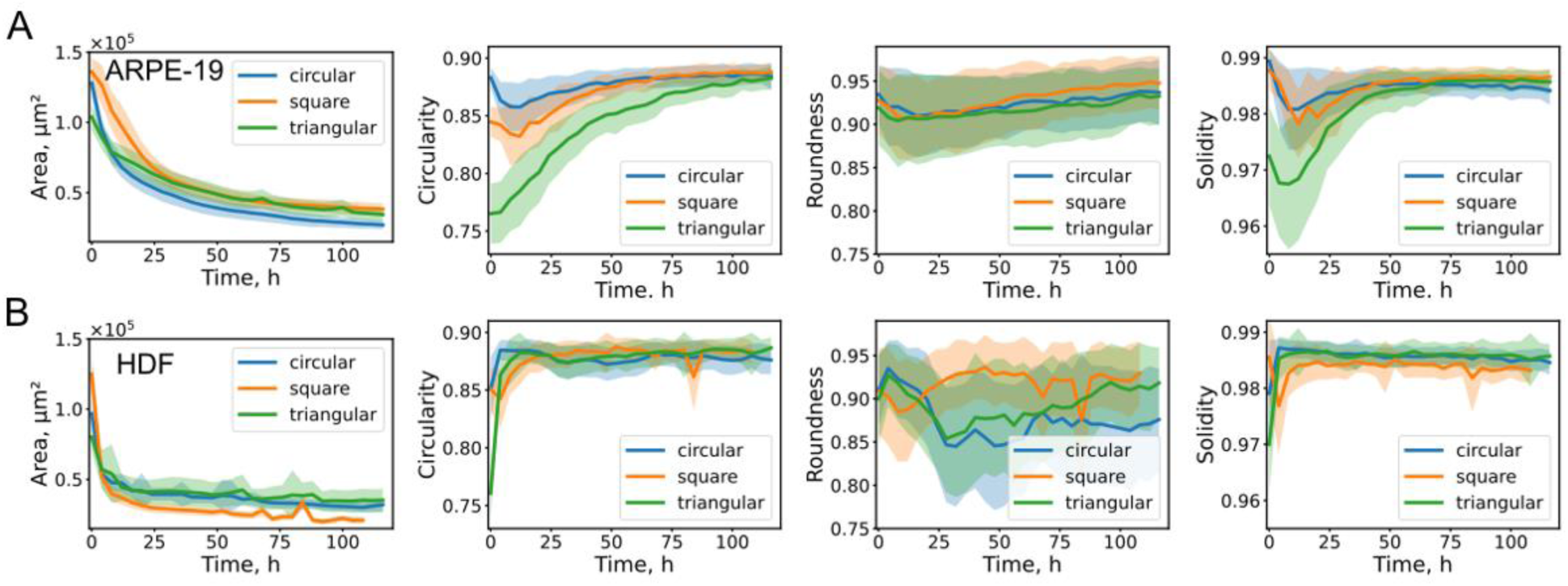
Time-dependent changes in morphological parameters of cell aggregates formed in circular, square, and triangular microwells, segmented using the SAMJ-IJ plugin. Data are presented separately for epithelial ARPE-19 cells (A) and mesenchymal HDF cells (B). The measured parameters include projected area (μm^2^), circularity, roundness, and solidity. Each curve represents the average across all spheroids per condition; shaded regions indicate the standard deviation.

Circularity describes how closely the object’s projection approximates a perfect circle (value of 1.0). At the initial time point, circularity was lowest in spheroids formed in triangular wells, due to the triangular cross-section being the furthest from a circle, followed by square wells (Fig. 3). Despite the compaction process, the final circularity value in all cases plateaued below 1.0 (typically around 0.9), reflecting minor irregularities in the fully formed spheroid outlines. Roundness, which indicates the degree of elongation, showed only modest changes over time, as the microwells lacked preferential axes of elongation (Fig. 3). Solidity, which is defined as the ratio of area to convex hull area, was initially low, reflecting irregular or loosely packed aggregate boundaries, and increased over time as the outer cell layers became more compact and cohesive (Fig. 3). A transient initial dip in these morphological parameters was observed, likely corresponding to the early cell adhesion phase and the morphological transition of the cell aggregates from the smooth microwell-defined shapes to actively reconfiguring spheroids.

Consistent with the faster compaction observed in HDF aggregates, both circularity and projected area values in square and triangular microwells reached their plateau levels significantly earlier compared to ARPE-19 aggregates. The time-dependent changes in these parameters were well described by exponential models: exponential rise to maximum for circularity and exponential decay for projected area, with a high goodness of fit (*R^2^* > 0.9). The characteristic time constant obtained from these fits was similar for both models and between the different types of microwells (Table 1), approximately four times lower for HDF cells (∼3–5 h) compared to ARPE-19 cells (∼22–28 h). This quantitative difference further supports the notion of cell-type-specific visco-capillary behavior influencing the kinetics of spheroid morphogenesis.

**Table 1.** Characteristic transition times and visco-capillary velocities estimated from projected area and circularity during spheroid formation in different types of microwells (circular, square, triangular). Values are reported as median ± half-width of the 95% confidence interval determined by bootstrap resampling. *n* indicates the number of spheroids analyzed across two biological replicates.

| Spheroid type | Type of microwells | $T_{\text{area}}$, h | $T_{\text{circ}}$, h | $v_p$ from $T_{\text{area}}$, $\mu\text{m/s}$ | $v_p$ from $T_{\text{circ}}$, $\mu\text{m/s}$ |
| --- | --- | --- | --- | --- | --- |
| ARPE-19 | circular (n=120) | 24±1 | - | 11±1 | - |
| | square (n=103) | 19±1 | 22±4 | 15±1 | 14±3 |
| | triangular (n=141) | 30±2 | 32±1 | 10±1 | 10±1 |
| HDF | circular (n=156) | 3.4±0.8 | - | 78±18 | - |
| | square (n=152) | 5.3±0.7 | 3.9±0.4 | 56±7 | 82±8 |
| | triangular (n=98) | 8.9±1.6 | 4.6±0.5 | 32±6 | 70±8 |

### Modeling the spheroid formation process

Spheroid formation, like many tissue organization processes, is widely accepted to resemble the behavior of liquids [22,42,43]. Based on this analogy, we employed a computational fluid dynamics (CFD) approach using the Volume of Fluid (VoF) method to simulate the morphogenesis of cell aggregates. The VoF method is an interface-capturing technique designed for simulating two immiscible fluid phases. In our model, one phase represented the cell aggregate, while the other corresponded to the surrounding cell culture medium.

Initial geometries were constructed to mimic the shapes imposed by the different microwell cross-sections (circular, square, and triangular) (Fig. 4, Movies [S4](https://drive.google.com/file/d/1ExPr4xZnI5zol2ZofKwz9Gp8BDTptTAR/view), [S5](https://drive.google.com/file/d/1sANs1Vkr4kLt4G3UqdN_WaS68iY9EvCt/view), [S6](https://drive.google.com/file/d/1JEP4i0LFfHLgBHpshupaazchk-QGd2uc/view)). The third dimension (thickness) of each initial shape was estimated based on the final spheroid geometry observed in experiments, assuming a constant volume, a final spherical shape, and using the average diameters from microscopic images. The estimated thickness was approximately 50 μm, which is smaller than the lateral dimensions (e.g., 200 μm for square microwells) due to both the number of seeded cells (2,000 per microwell) and the microwell area **used**.

**Figure 4.**
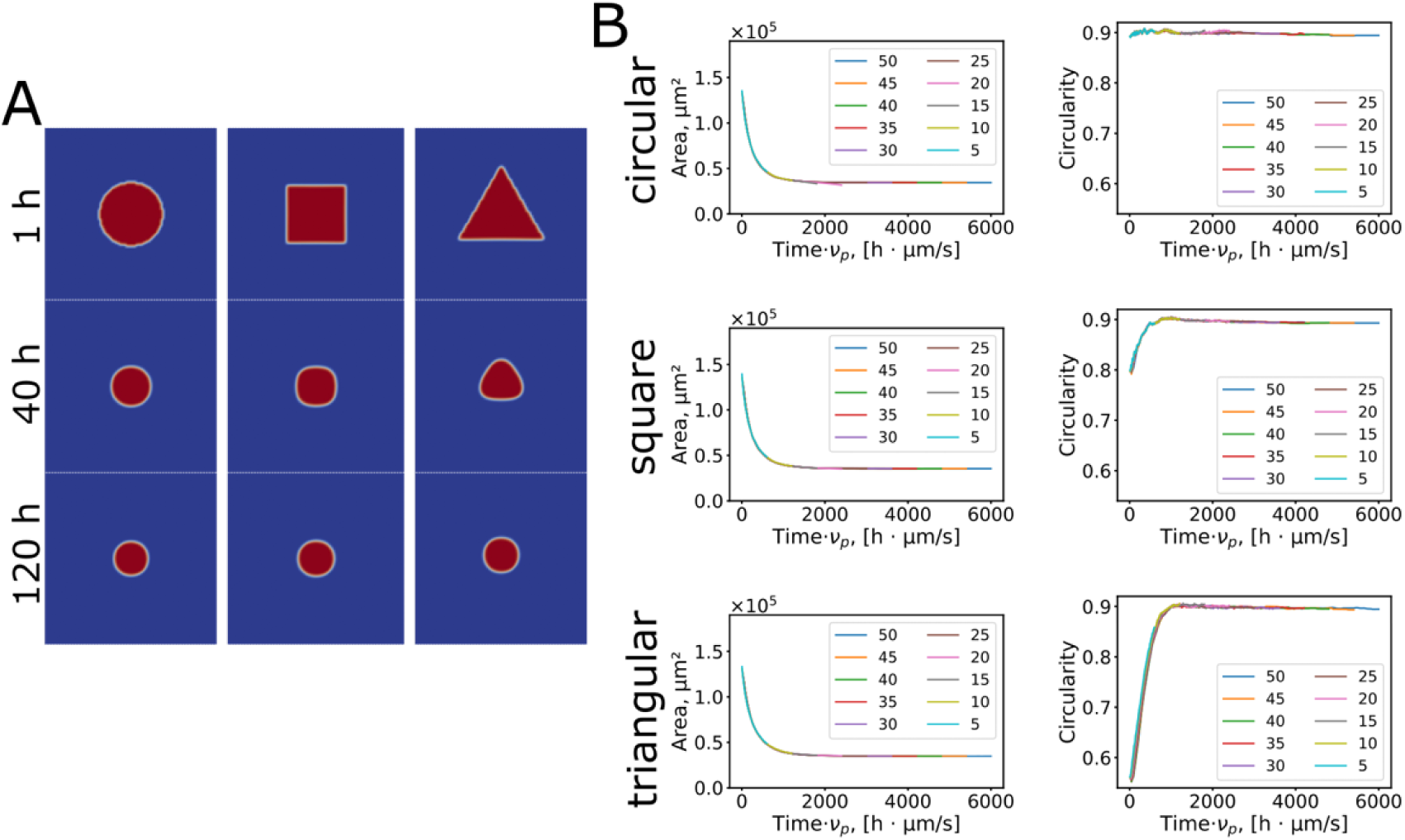
Computational fluid dynamics (CFD) modeling of cell aggregation dynamics. **(A)** Simulated horizontal projections of spheroid compaction within circular, square, and triangular microwells **at** a visco-capillary velocity of 15 μm/s. Snapshots are shown at 1 h, 40 h, and 120 h to illustrate the morphological evolution over time. **(B)** Normalized changes in projected area (μm²) and circularity as a function of normalized time (defined as time multiplied by visco-capillary velocity) for a range of visco-capillary velocities (5–50 μm/s). The curves illustrate how the compaction dynamics scale consistently with the visco-capillary parameter across different geometries.

Simulations were run over time intervals matching the experimental observations (120 h total). Values for surface tension (σ) and viscosity (η) were assigned to the cell phase to match the experimental dynamics. Across all geometries, the simulations showed a gradual morphological transition from the initial shapes (cylindrical, cuboidal, or triangular prismatic forms) to nearly spherical aggregates (Fig. 4, Movies [S4](https://drive.google.com/file/d/1ExPr4xZnI5zol2ZofKwz9Gp8BDTptTAR/view), [S5](https://drive.google.com/file/d/1sANs1Vkr4kLt4G3UqdN_WaS68iY9EvCt/view), [S6](https://drive.google.com/file/d/1JEP4i0LFfHLgBHpshupaazchk-QGd2uc/view)). The speed of this transformation was governed by the ratio of surface tension (σ) to viscosity (η), known as the visco-capillary velocity (*ν_p_ =* σ/η).

When the simulation time was scaled by the visco-capillary velocity (i.e., multiplied by this parameter), the resulting dynamics overlapped, showing identical behavior for all visco-capillary velocity values within a given initial geometry. This outcome aligns with theoretical expectations based on fluid mechanics. In highly viscous systems such as those simulated here, inertial forces are negligible, and the unsteady Navier–Stokes equations reduce to a linear form with the nonlinear inertial term omitted. As a result, the characteristic time scale of the system exhibits inverse scaling with respect to both material and geometric parameters. If the ratio σ/η is scaled by a factor X, the characteristic time t must scale inversely as t ∝ 1/X. Similarly, if all spatial dimensions are divided by a factor X, then t ∝ 1/X as well [44].

The parameters circularity and projected area (excluding those from circular microwells) exhibited exponential rise and decay trends, respectively, closely mirroring the experimental observations (Fig. 5, [Movie S7](https://drive.google.com/file/d/1eNIhOaNTSM_mrMpKrZ6BDU7T1nEP7C8o/view)). Minor discrepancies between simulated and experimental values, particularly at the initial and final stages of circularity, were attributed to greater morphological irregularities and surface roughness in real spheroids. The decrease in projected area was similar between experiment and simulation, and also highly consistent across all microwell geometries, suggesting that its dynamics are governed primarily by the initial width-to-thickness ratio of the cell layer, rather than by the specific cross-sectional shape of the microwell (Fig. 4B).

**Figure 5.**
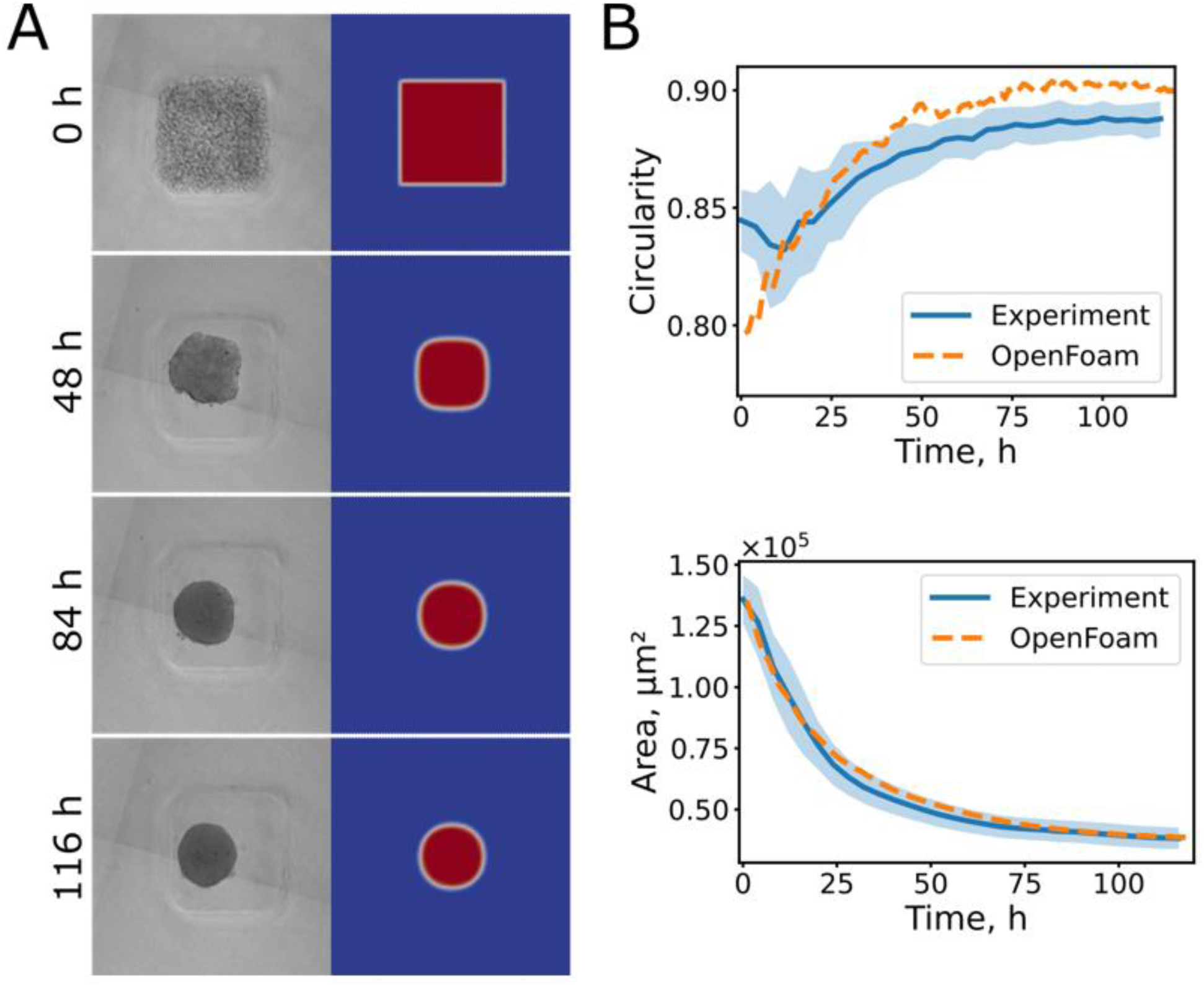
Direct comparison of experimental spheroid formation data and OpenFOAM simulation results. A representative case of ARPE-19 cell aggregation in square microwells; OpenFOAM simulation with the same geometric parameters and a visco-capillary velocity of 10 μm/s. (A) Side-by-side snapshots showing microscopy observations of spheroid formation and corresponding OpenFOAM simulation outcomes at four time points (0 h, 48 h, 84 h, and 116 h). (B) Extracted plots of circularity and projected area (μm²) as a function of time from both experimental measurements (solid curve; mean of 46 spheroids formed in microwells within the same well; shaded region indicates standard deviation) and simulations (dashed line).

By fitting exponential functions to the simulated datasets, we determined characteristic transition times for each microwell geometry. Transition times derived from the projected area decay (τ_area_) were consistent across geometries: 264±3, 295±3 and 288±3 h·μm/s for circular, square, and triangular microwells, respectively (values scaled by the visco-capillary velocity; errors represent the fitting uncertainty). Transition times calculated from circularity dynamics (τ_circ_) were also in close agreement, with a value of 320±3 h·μm/s for both square and triangular microwells. These results indicate that the two approaches, which are based on projected area and circularity, yield similar time constants. However, discrepancies may arise under different cell seeding densities, as the projected area is more sensitive to initial cell volume (i.e., the width-to-thickness ratio). The extracted time constants allowed alignment between simulated and experimental curves for both circularity and projected area across all microwell shapes, as shown for a representative example of ARPE-19 spheroids in square microwells in Fig. 5. This allowed us to estimate the visco-capillary velocity for a given well geometry by comparing the characteristic transition times between simulation and experiment.

### Estimating the visco-capillary velocity based on the spheroid formation process

The extracted transition times for ARPE-19 and HDF spheroids in different types of microwells are presented in Table 1. Calculated visco-capillary velocities were quite consistent across microwell types and between the two evaluation approaches (projected area and circularity): 10**–**15 μm/s for ARPE-19 spheroids and 30–80 μm/s for HDF spheroids (averaged across all measurement approaches). The latter exhibited an approximately fivefold faster compaction rate, in agreement with previous morphological observations.

Using the experimentally determined visco-capillary velocity, either surface tension or effective viscosity can be estimated, provided that one of these parameters is independently measured. In this study, additional mechanical characterization was performed using atomic force microscopy (AFM) and parallel-plate compression to determine the surface tension of ARPE-19 and HDF spheroids (details are provided in the Supplementary Material). As described in our previous work (see Supplementary Information and [35]), this approach combines measurements of local stiffness (AFM) and global mechanical response (compression), enabling estimation of surface tension from its differential contribution to the two measurements (Fig. S4, Table S1). The model treats the spheroid as an effectively elastic body with intrinsic surface tension arising from active contractile forces generated by myosin motors in the actin cortex of the cells in the spheroid’s outer layer. At the short timescale of the AFM and parallel-plate compression measurements (on the order of seconds), contributions from active cortex remodeling and poroelastic flow are expected to be minimal, so the simplified model provides a reasonable approximation of the mechanical response. The resulting surface tension values were 0.28 ± 0.10 N/m for ARPE-19 spheroids and 1.1 ± 0.2 N/m for HDF spheroids. Using these values together with the corresponding visco-capillary velocities (derived from τ_circ_ in square microwells for consistency), we estimated the effective viscosity η of the aggregates as (5.0±0.1)·10^4^ Pa·s for ARPE-19 and (7.3±0.1)·10^4^ Pa·s for HDF spheroids. These results indicate that the observed differences in visco-capillary velocity between the two cell types are primarily driven by differences in surface tension, while HDF spheroids also exhibited a moderately higher effective viscosity.

In principle, both the time dependence of circularity and projected area can be used to determine the visco-capillary velocity using the proposed approach. However, circularity is informative only for non-circular microwells, as the initial configuration in circular wells already exhibits high circularity, thereby limiting its dynamic range. In contrast, changes in projected area are applicable to all microwell geometries but are strongly influenced by the initial cell volume (i.e., the thickness of the cell layer within the well and the seeding density), since the reduction in projected area is primarily driven by vertical redistribution of cells during compaction (Movie [S5](https://drive.google.com/file/d/1sANs1Vkr4kLt4G3UqdN_WaS68iY9EvCt/view)). For example, when the initial cell layer is thick (high cell number within a small well area), the projected area may change only minimally over time. This behavior was also reproduced in simulations by systematically increasing the width-to-thickness ratio (Fig. 6A). With increasing ratio, the decrease in projected area became progressively slower, and at a width-to-thickness ratio of 0.75, the projected area even increased over time. In contrast, the dynamics of circularity were substantially more robust to variations in seeding density. Similar characteristic times (variation ∼10%) were obtained across width-to-thickness ratios ranging from 0.05 to 0.75. Moreover, circularity-based analysis yielded consistent characteristic times across different microwell geometries (Fig. 6B), indicating that circularity provides a more stable descriptor for extracting visco-capillary kinetics.

**Figure 6.**
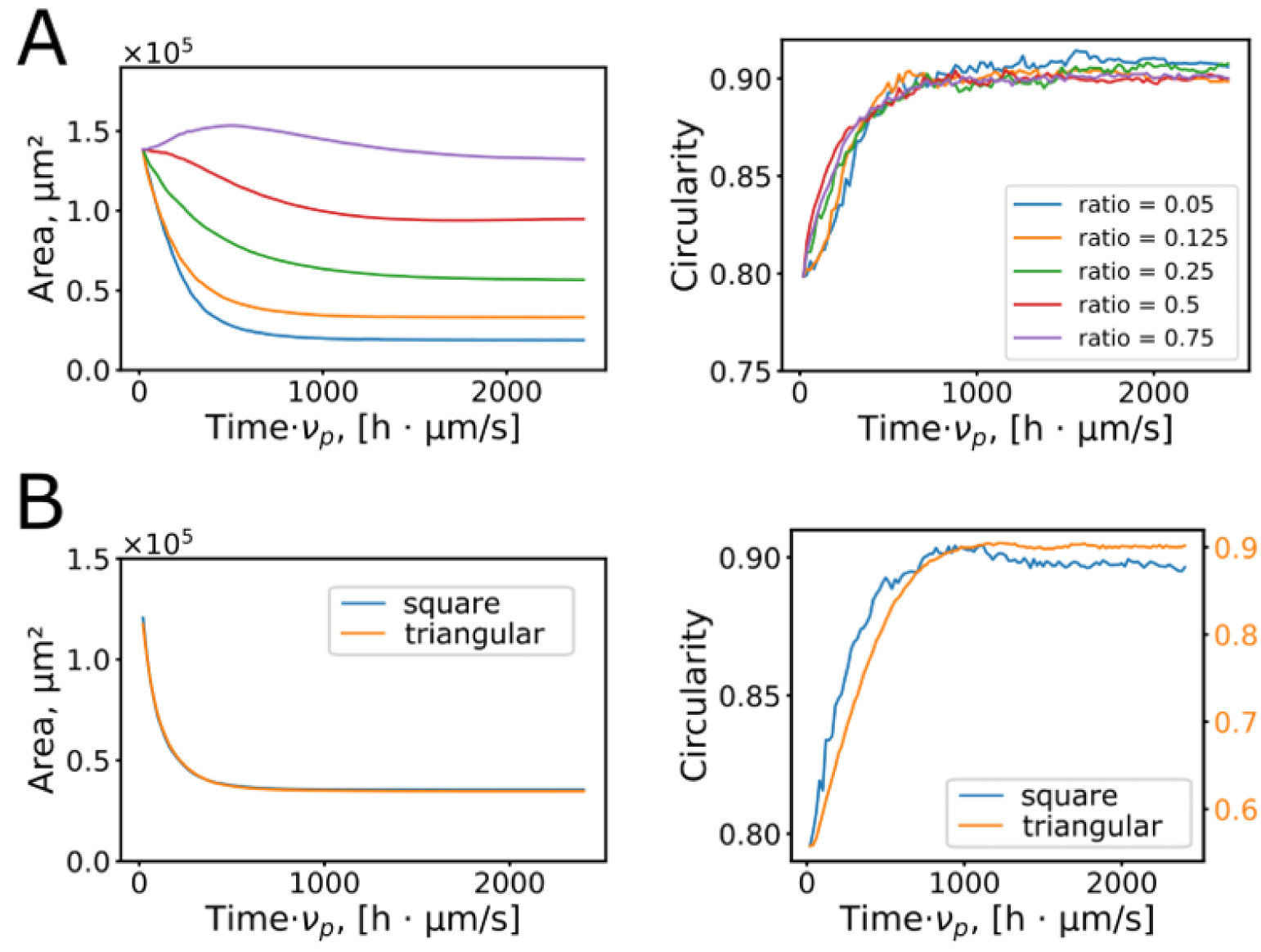
Effects of width-to-thickness ratio (cell seeding density) and microwell cross-sectional geometry on the dynamics of projected area and circularity. (A) The dynamics of the projected area are highly sensitive to the width-to-thickness ratio of the initial cell layer (i.e., seeding density). At high ratios, the projected area changes only slightly or even increases over time, while the dynamics of circularity remain stable. (B) When the width-to-thickness ratio is held constant (here, 1/8), the projected area dynamics remain consistent across different microwell geometries (square and triangular). Circularity demonstrates a similar characteristic time in both geometries, although the starting values differ (note the differing scales and starting values between parameters).

Therefore, the optimal experimental setup for estimating visco-capillary velocity is to track the dynamic change in circularity within square microwells. Square wells provide a well-defined initial geometry while avoiding the limitations associated with circular wells (high initial circularity). Triangular microwells were found to be less reliable due to the presence of sharp corners, which are not always accurately reproduced during 3D printing and tend to become rounded during agarose casting. Furthermore, cells often avoid the corner regions of the triangular wells, resulting in an initial aggregate configuration that deviates from the ideal triangular prism shape (Fig. S3). These deviations introduce inconsistencies in the compaction dynamics and reduce the reliability of morphological tracking in this geometry.

### Modulating spheroid formation and shape engineering

To investigate the role of cellular contractility and surface tension in spheroid morphogenesis, we applied a set of pharmacological treatments designed to modulate cytoskeletal dynamics and effective surface tension. These included blebbistatin (a myosin II inhibitor), Y-27632 (a Rho-associated kinase inhibitor), and cytochalasin D (cytD), which disrupts actin polymerization. The agents were added right after seeding the cells into the microwells and were present throughout the entire spheroid formation time. Transition times and visco-capillary velocities were estimated from circularity changes during spheroid formation (Fig. 7, Movie [S8](https://drive.google.com/file/d/1jEy_qU4ZX_u9hIyy2VZT9WDO2AtueJRM/view), Movie [S9](https://drive.google.com/file/d/1IPHDD51bRDnszMrBNyNkwd2bjrOBTDMY/view)).

**Figure 7.**
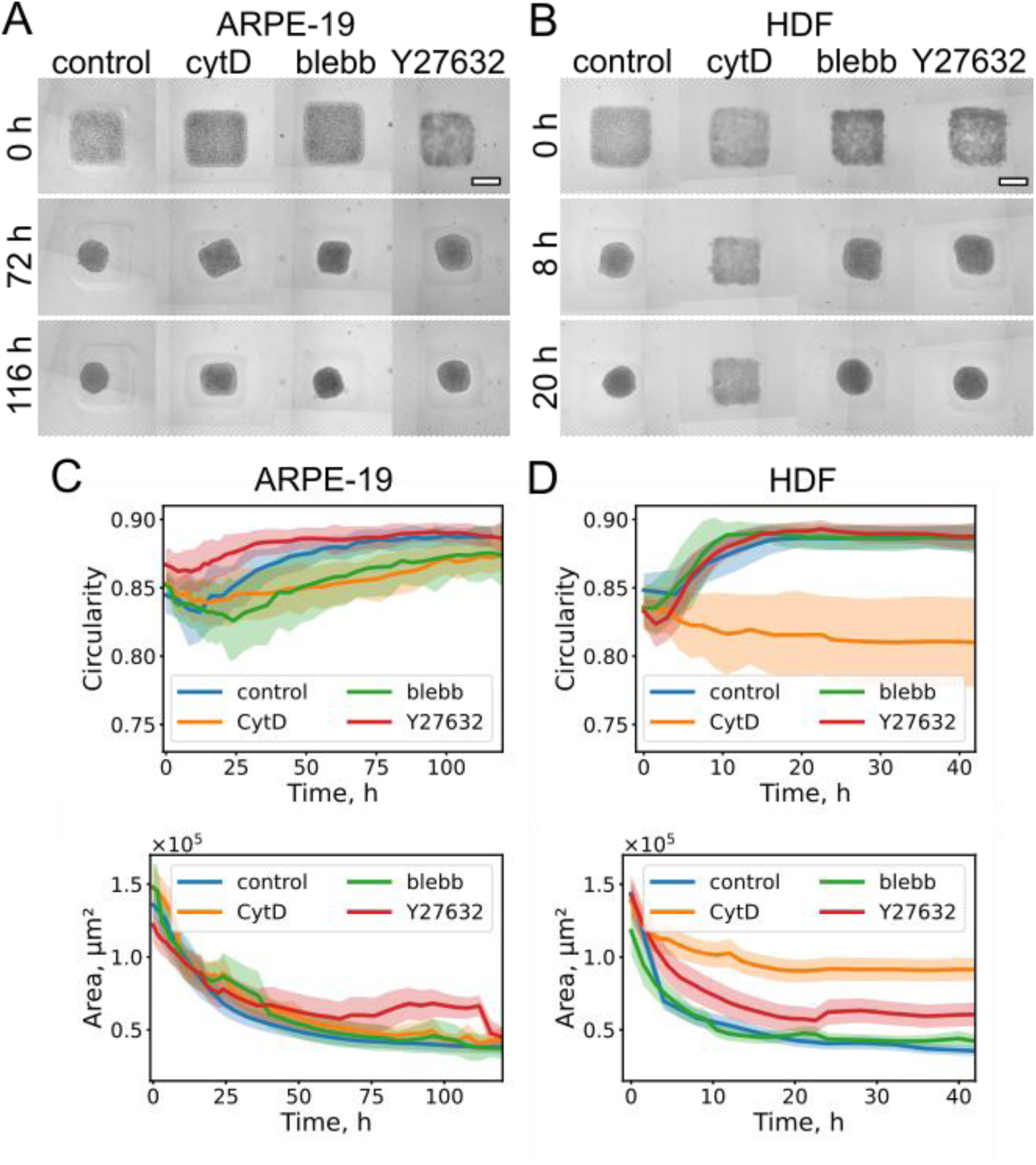
Modulation of the spheroid formation process by different treatments affecting surface tension (cytochalasin D (cytD), blebbistatin (blebb), and Y-27632). (A, B) Time-resolved observation of spheroid formation from ARPE-19 (A) and HDF cells (B) in square microwells at the indicated time points. Scale bar: 200 μm. (C, D) Corresponding changes in circularity and projected area.

Blebbistatin and cytD treatments significantly increased transition times in both ARPE-19 and HDF spheroids (Table 2), while Y-27632 slowed formation only in HDF spheroids (ASL<0.01 from a nonparametric permutation test). Shape descriptors at 100 h and 20 h for ARPE-19 and HDF spheroids, respectively, were analyzed (Figs. S5, S6, Table S2). Among all shape descriptors, the most pronounced changes were observed as a decrease in circularity and an increase in curvature variance (Var(κ)) and characteristic maximum curvature (κ_max_) for cytD-treated spheroids.

**Table 2.** Transition times and visco-capillary velocities estimated from circularity during spheroid formation in square microwells with different treatments (cytochalasin D (cytD), blebbistatin, and Y-27632). Median ± half-width of the 95% confidence interval (CI) determined by bootstrapping, *n* – number of spheroids analyzed from two biological repeats.

| Spheroid type | Treatment | $T_{\text{circ}}$, h | $v_p$ from $T_{\text{circ}}$ $\mu\text{m/s}$ |
| --- | --- | --- | --- |
| ARPE-19 | control (n=135) | 35±5 | 9.1±1.3 |
| | cytD, 0.5 $\mu\text{M}$ (n=178) | >1000 | - |
| | blebbistatin, 10 $\mu\text{M}$ (n=178) | 92±31 | 3.5±1.2 |
| | Y-27632, 50 $\mu\text{M}$ (n=124) | 21±3 | 15±2 |
| HDF | control (n=158) | 5.3±0.7 | 60±8 |
| | cytD, 1 $\mu\text{M}$ (n=134) | >1000 | - |
| | blebbistatin, 10 $\mu\text{M}$ (n=172) | 16±3 | 20±4 |
| | Y-27632, 50 $\mu\text{M}$ (n=161) | 10±1 | 32±3 |

Therefore, cytD had the most pronounced effect on spheroid formation from both ARPE-19 and HDF cells among all applied treatments. At high concentrations (>2 μM), cytD completely blocked aggregation, while at lower concentrations (0.5 μM for ARPE-19, 1 μM for HDF), aggregates still formed and compacted but largely retained the geometry of the microwell (e.g., square cross-section) (Fig. 7, Movie [S10](https://drive.google.com/file/d/1x4w0fCMa0P2BSZvWOCnY2Kxvsfh9KvlO/view)). While both circularity and projected area changed more slowly than in control microwells, the effect on circularity was much more substantial. This behavior suggests a reduction in surface tension was sufficient to prevent the usual rounding transition characteristic of spheroid compaction. However, there was still cell adhesion and aggregate compaction, since the projected area decreased and the formed aggregates acquired shapes reflecting the shape of the initial microwell (square cross-section).

Since cytD strongly affects the actin cytoskeleton structure and dynamics, we performed additional experiments to verify whether these effects were reversible. First, cytD-treated spheroids were washed while still confined in microwells and monitored for shape recovery (Fig. 8B). Second, treated spheroids were extracted from the microwells and seeded onto standard tissue culture plastic to assess cell outgrowth (Fig. 8C). HDF spheroids gradually recovered their shape within 4–5 days, with a characteristic recovery time of 109 ± 15 h, which was substantially longer than the characteristic time of aggregation from suspension. Circularity returned to 0.88 ± 0.01, comparable to control spheroids. In contrast, ARPE-19 spheroids largely retained their square shape for at least 7 days, exhibiting circularity values of 0.86 ± 0.02 at day 7 compared to 0.85 ± 0.02 at the onset of recovery (p < 0.05). Despite the persistent macroscopic shape in ARPE-19 aggregates, seeding onto culture plastic induced active cell outgrowth (Fig. 8C), indicating preserved cytoskeletal dynamics and cell viability following cytD treatment. The area of cell outgrowth was quantified at day 7 for ARPE-19 spheroids and at day 4 for HDF spheroids. For ARPE-19 brick-like spheroids, the outgrowth area was only 16% lower than that of control spheroids (p < 0.05). In contrast, HDF spheroids showed a more pronounced reduction in outgrowth area (60% lower than control, p < 0.0001).

**Figure 8.**
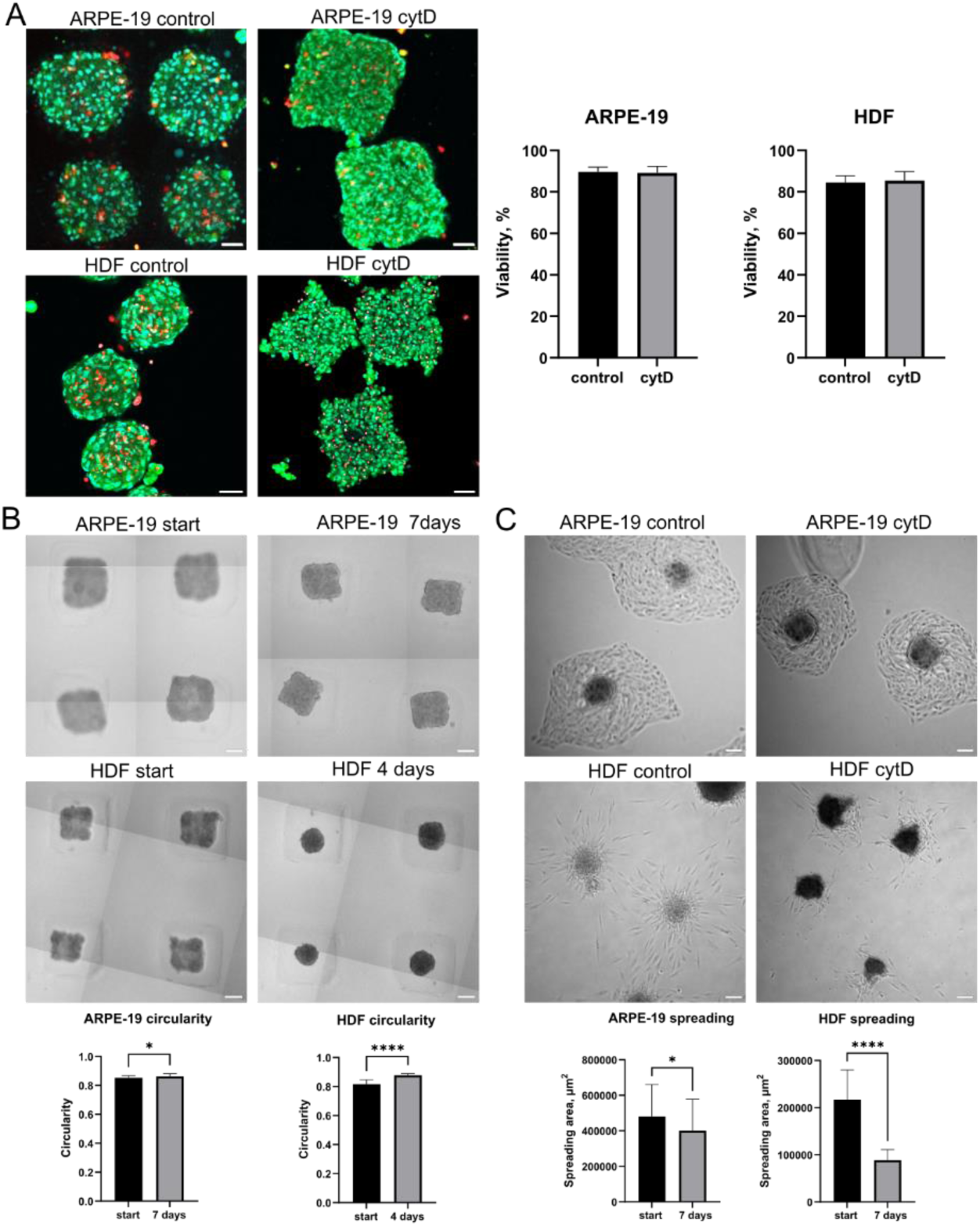
Engineering of spheroid shapes through the combination of microwell confinement and cytochalasin D (cytD) treatment. (A) Viability of control and cytD-treated ARPE-19 spheroids generated in square microwells. Scale bar is 50 μm. (B) Square ARPE-19 and HDF spheroids after cytD washout, shown at the onset of recovery and after several days (7 days for ARPE-19; 4 days for HDF). Scale bars are 100 μm. Bottom: quantified circularity. (C) Control and square cytD-treated ARPE-19 and HDF spheroids after cytD washout and subsequent seeding onto cell culture dishes (7 days for ARPE-19; 4 days for HDF). Scale bars are 100 μm. Bottom: quantified spreading area.

Control spherical spheroids and cytD-induced brick-like spheroids were fixed and stained to visualize the actin cytoskeleton and tight junctions in ARPE-19 spheroids using immunostaining for the tight junction protein ZO-1. As expected, cytD treatment induced marked alterations in the actin cytoskeleton (Fig. 9A,B), resulting in a less developed and more disrupted fibrillar network compared to control spheroids. Confocal Z-stacks were acquired to quantify spheroid height. CytD-treated spheroids exhibited significantly reduced heights compared to controls (ARPE-19: 43 ± 11 μm, n = 16, vs. 111 ± 25 μm, n = 11; HDF: 57 ± 21 μm, n = 30, vs. 106 ± 17 μm, n = 16; Fig. 9D). Tight junctions remained detectable in ARPE-19 spheroids following cytD treatment, although their organization appeared less developed compared to that in control spheroids (Fig. 9C).

**Figure 9.**
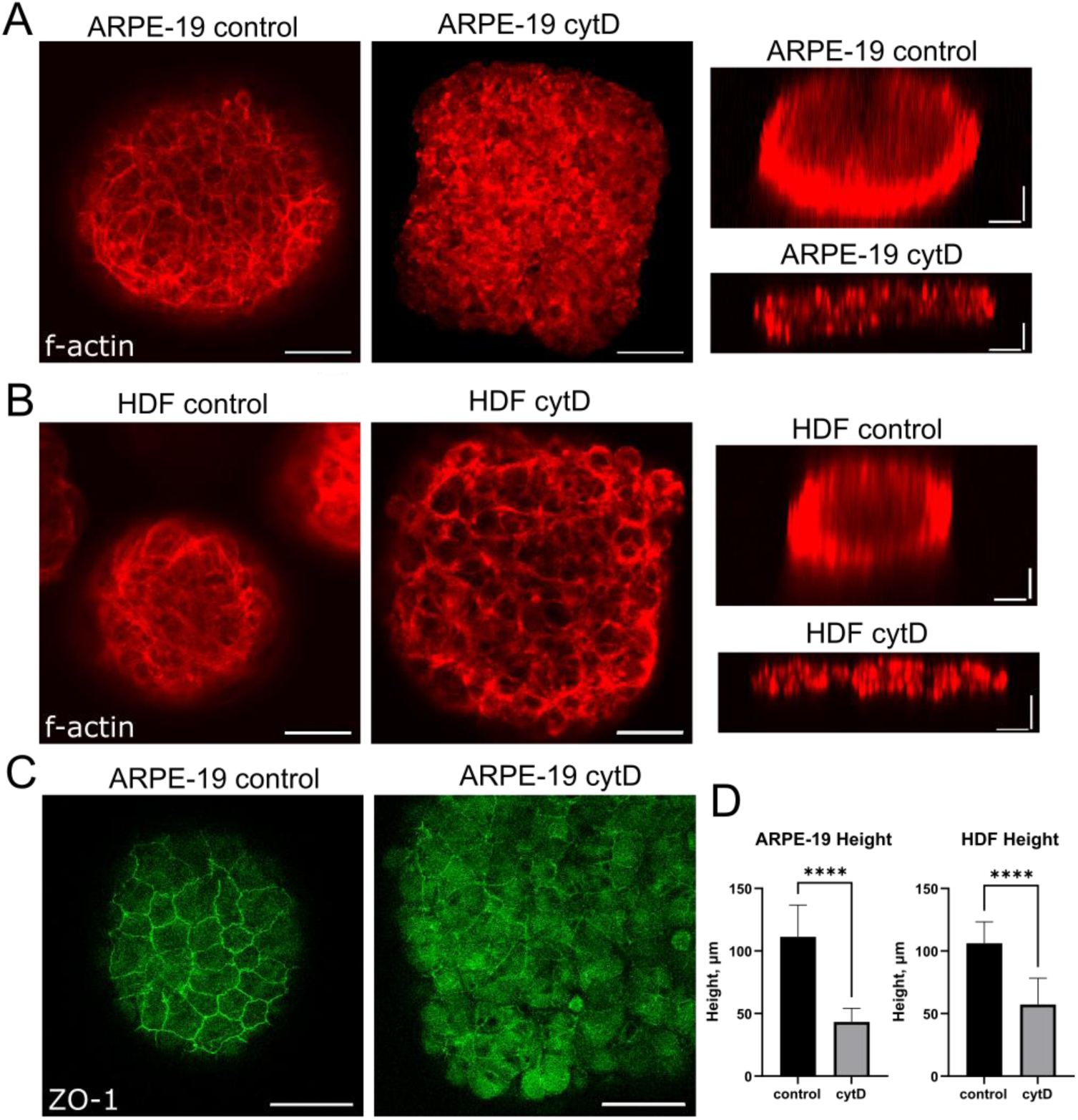
Confocal microscopy of fixed and stained control and cytD-treated square ARPE-19 and HDF spheroids. (A) F-actin staining in ARPE-19 spheroids. Scale bars are 50 μm. (B) F-actin staining in HDF spheroids. Scale bars are 50 μm. Right panels: vertical cross-sections from Z-stacks. Scale bars are 30 μm (both horizontal and vertical). (C) Tight junction staining in ARPE-19 spheroids using ZO-1 antibodies. (D) Quantification of spheroid height from Z-stacks of F-actin staining.

To further demonstrate the potential of this shape-engineering approach, microwells with a more complex star-shaped cross-section were fabricated. Under control conditions, cell aggregates adopted a typical spherical morphology, whereas in the presence of cytD, distinct star-shaped spheroids were formed (Fig. 10A, Movie [S11](https://drive.google.com/file/d/1BruQiGDoa8QH5p4kgLRec1x_IheuW9gJ/view)). These spheroids were successfully extracted from the microwells and retained their characteristic geometry after removal. For all spheroid types, local boundary curvature and global shape descriptors were quantified (Fig. 10B, C, Fig. S7, Table S3). Among the engineered geometries, triangular and star-shaped spheroids showed the greatest deviation from control spherical aggregates, with lower circularity and a higher perimeter-to-normalized-area ratio.

**Figure 10.**
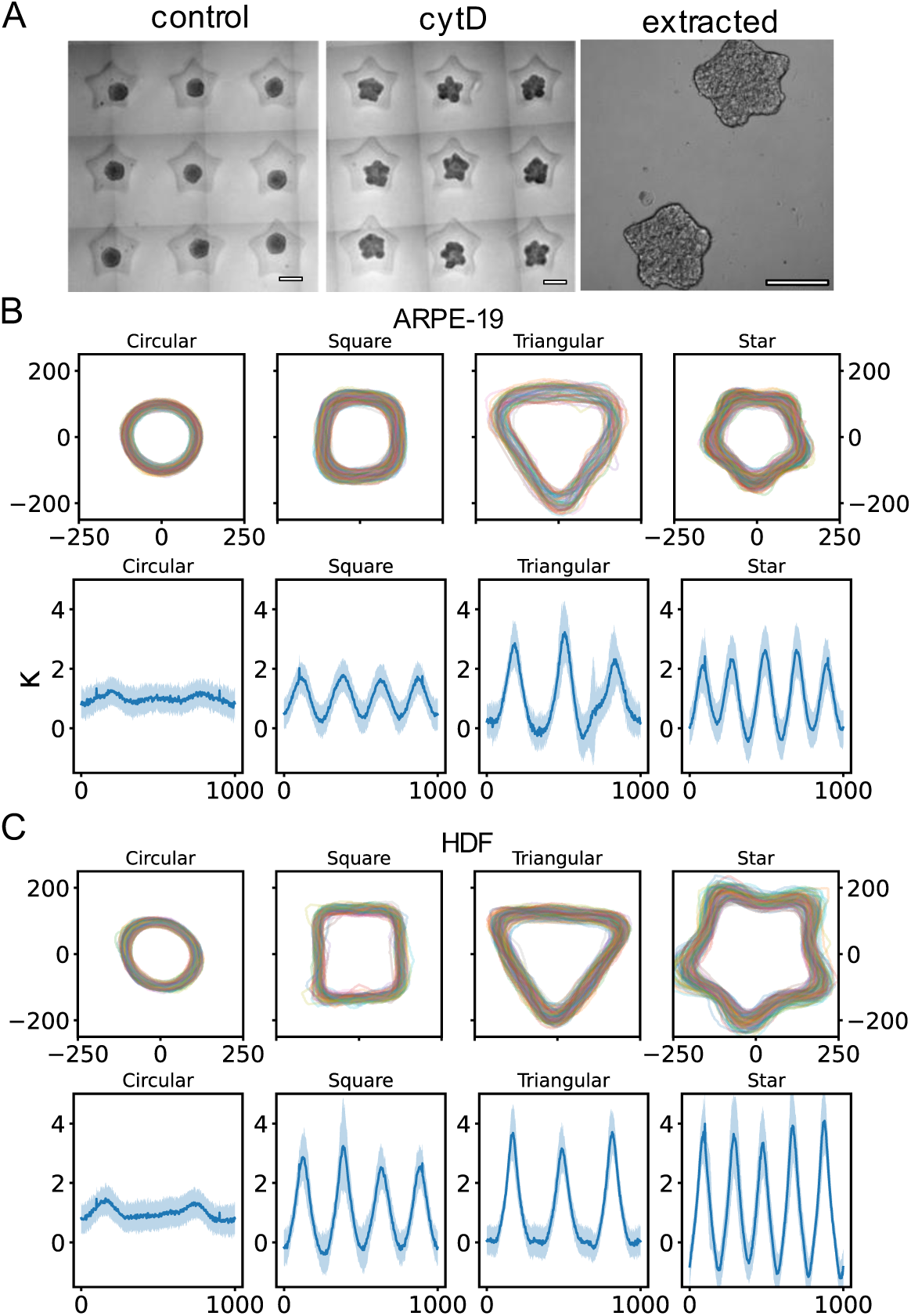
Morphology of star-shaped and other shape-engineered spheroids. (A) Generation of star-shaped spheroids. Left: control aggregates; middle: CytD-treated aggregates acquiring star-shaped morphology within the corresponding microwells; right: extracted spheroids retaining their star-shaped structure. Scale bars: 200 μm. (B) Aligned contours and corresponding local boundary curvature profiles for all types of shape-engineered ARPE-19 spheroids (circular, brick-like, prismatic, and star-shaped). (C) Aligned contours and corresponding local boundary curvature profiles for all types of shape-engineered HDF spheroids (circular, brick-like, prismatic, and star-shaped).

Analysis of local curvature profiles revealed that the number of prominent curvature peaks corresponded to the imposed geometry (four for square, three for triangular, and five for star-shaped spheroids), indicating preservation of angular features (Fig. 10). Consistently, both curvature variance (Var(κ)) and the characteristic maximum curvature (κ_max_) were significantly higher in shape-engineered spheroids compared to controls, quantitatively confirming partially preserved corner sharpness and deviation from circular geometry.

Due to their larger projected perimeter and reduced height (thickness), as determined from confocal Z-stacks, shape-engineered spheroids exhibited a substantially higher surface-to-volume ratio (up to twofold) compared to conventional spherical spheroids (see Supplementary Information for calculations). Importantly, this increase was achieved while maintaining a comparable overall lateral size and aspect ratio. The elevated surface-to-volume ratio is expected to enhance mass transport across the aggregate boundary, facilitating improved diffusion of oxygen and nutrients as well as the removal of metabolic waste.

## Discussion

Here, we propose an approach to estimate visco-capillary velocity during spheroid formation in microwells of defined shapes based on temporal changes in either projected area or circularity. The projected area-based approach can be applied to standard circular wells, but it requires careful control of cell seeding density and estimation of the initial cell layer thickness. In our case, thickness was inferred from the final spheroid volume under the assumption of volume conservation. This assumption may not hold for rapidly proliferating spheroids, such as those formed by transformed or cancer cell lines [45–47], where progressive increases in all aggregate dimensions are expected over time. For primary and non-transformed cells, aggregate volume may even decrease over time due to cellular reorganization, elimination of intercellular cavities, or even reduction in cell size [45,47–49]. Indeed, in our experiments, a two-stage decrease in projected area was observed, with a rapid initial compaction followed by a slower stabilization phase (Fig. 3). Circularity, on the other hand, is more robust to variations in cell number, but requires the use of non-circular microwells. Although such wells are not yet common, advances in 3D printing make their fabrication relatively straightforward [39,41]. In our setup, visco-capillary velocities calculated from both projected area and circularity were consistent when the CFD model was parameterized for the experimentally determined aggregate volume.

Image analysis has proven to be a valuable tool for the quantitative assessment of 3D cell culture dynamics [46,47,50–54]. High-content imaging combined with automated segmentation enables precise tracking of morphological parameters such as spheroid size, circularity, and surface irregularities over time, thereby providing high statistical power. Recently, AI-based approaches for spheroid segmentation have gained popularity [54–56], and their implementation in open-source software has simplified their application for end users [57]. The integration of automated image analysis with computational modeling, as conducted here, offers a powerful route for linking experimental observations to biophysical parameters, with visco-capillary velocity being one such parameter.

The idea of describing the dynamics of cell populations in terms of a viscous liquid model was developed in the seminal works of Steinberg et al. [22,23,58,59] and has since been quite fruitful in the description of cell sorting, aggregate fusion, spreading, and other phenomena [21,27,32,60,61]. A formal analogy with the rheology of viscous liquids was suggested, where restructuring processes are driven by surface tensions and opposed by cell/tissue viscosity. In particular, tissue viscosities were estimated by comparing the dynamics of cell aggregates with the dynamics of viscous liquids placed in similar initial geometrical configurations [23,44]. It was suggested and confirmed that rounding of cell aggregates follows an exponential dependence and depends on surface tension and aggregate size [44].

While multicellular spheroids are considered viscoelastic, elasto-visco-plastic, or more complex systems [62], in the present work, the modeling was focused specifically on the early-stage rounding and compaction dynamics of cell aggregates occurring over timescales of tens of hours, during which the system behaves effectively as a highly viscous fluid. The use of a Newtonian fluid with surface tension in the CFD model represents an effective description of the long-time viscous limit of spheroid mechanics. Previous experimental studies [35,63] and the present study as well have shown that cell spheroids exhibit stress relaxation on characteristic timescales ranging from seconds to minutes, depending on cell type and experimental setup (here, 2.7 ± 0.8 s for ARPE-19 and 4.3 ± 1.1 s for HDF spheroids in parallel-plate compression experiments). Assuming a characteristic relaxation time of the aggregates on the order of τ_rel_∼0.01 - 0.1 h, and a characteristic spheroid formation time of τ∼10–100 h, the Deborah number De= τ_rel_ / τ is in the range De∼10^−4^–10^−2^, supporting the fluid-like approximation on the timescales considered here. Similarly, the effective Weissenberg number, defined as the shear rate times the relaxation time, remains well below unity due to the extremely low strain rates associated with spheroid rounding. These estimates indicate that elastic effects are likely negligible during the observed dynamics.

Prior works utilized spontaneously or specially formed elliptical cell aggregates which then rounded into a sphere, but without the support of corresponding geometric computational models [44,64]. Such approaches are less controlled and reproducible than the use of defined microwell geometries applied in the present study. Another advantage of the proposed approach is that, due to the small dimensions of the microwells (comparable to those of conventional round microwell systems), a large number of spheroids can be generated simultaneously, enabling their use in subsequent experiments.

On the other hand, several previous studies have demonstrated deviations from the viscous liquid model in the behavior of cell aggregates [17,27,65]. Some of these deviations can be attributed to the accumulation of ECM within spheroids over time [47,66,67]. In this regard, the present analysis of spheroid formation reflects an early stage, when ECM deposition is still minimal, since ECM accumulation typically occurs over several days to weeks [47,66]. Consequently, the estimations obtained here are closer to the intrinsic behavior of cells that are not yet restricted by ECM. These results can therefore be compared with experiments on fully matured spheroids to disentangle the contribution of ECM. In this context, the visco-capillary velocities estimated in our study (∼10–100 μm/s) are notably higher than those typically reported for fusion of mature spheroids (∼10–100 μm/h) [21,27]. The slower kinetics observed in spheroid fusion and in the recovery of square aggregates toward a spherical shape may be attributed to the presence of ECM and pre-established cell–cell contacts, which can increase mechanical resistance and promote a jamming-like state [65,68]. Such structural maturation is expected to reduce effective fluidity compared to newly formed aggregates. However, further experiments are required to directly confirm this mechanism and to disentangle it from potential contributions of cytD treatment applied here.

CFD approaches for modelling multicellular aggregates have been used previously in several works. In particular, phase-field methods have been applied to model the formation, fusion, and morphodynamics of cellular spheroids in the context of tissue biofabrication and mechanobiology [32,69,70]. For instance, Yang et al. employed a phase-field model to simulate the fusion of cellular aggregates embedded in a hydrogel, showing that tissue spheroids behave similarly to viscous droplets with tunable surface tension and viscosity parameters [32]. Similarly, Cristea and Neagu applied the lattice Boltzmann method to study shape changes of tissue constructs, reinforcing the analogy between multicellular spheroids and soft viscoelastic materials [69]. More recently, Sciumè et al. introduced a Cahn–Hilliard–Navier–Stokes phase-field model to quantify the rheological behavior of soft cell aggregates under micropipette aspiration, offering insights into interfacial tension and aggregate deformation [71]. Lin et al. also demonstrated the use of an active phase-field model to simulate gut spheroid morphogenesis, incorporating feedback between mechanical instabilities and actomyosin contractility [72].

Collectively, these studies highlight the versatility of phase-field models in capturing both the mechanical and morphological aspects of spheroid dynamics. While the VoF approach has been widely used in engineering fluid simulations, its application to biological systems, especially to model tissue morphogenesis and spheroid compaction, is relatively novel. The VoF method, used in our study, offers a more direct and mass-conservative approach to modeling sharp fluid–aggregate boundaries, and its application to spheroid rounding kinetics represents a novel contribution to the field. An additional advantage is that the VoF method is available in the open-source CFD software package OpenFOAM. The model can be easily extended to viscoelastic formulations, such as the Oldroyd-B model [73], which are already implemented in OpenFOAM. The obtained results confirm that the dynamics of spheroid formation follow scaling laws governed by the visco-capillary ratio, reinforcing the physical validity of using a VoF-based fluid model to describe cell aggregate behavior.

In the current modeling setup, we have neglected the interactions of the cells with the microwell walls. However, this simplification is reasonable, since, starting from the initial moment, the projected area of the cell aggregate decreases and cells lose contact with the side walls. When the number of cells is sufficiently large (i.e., when the thickness of the cell layer is large), the shape transition will be constrained by the walls, and the maximum projection dimensions and area will be limited by the microwell shape.

While spherical spheroids remain the dominant variant used, due to the common use of circular wells and the action of surface tension during cellular aggregation, the current study provides a way to explore the potential of non-spherical geometries, such as brick-like cell spheroids, as novel building blocks for tissue engineering and the creation of complex multicellular architectures. Ultrahigh cell-density 3D bioprinting via acoustic-fluids-mediated stereolithography was recently developed, enabling the fabrication of arbitrary multicellular architectures [74]. In another study, engineered 2D and quasi-3D extracellular matrices were used to demonstrate how interfacial geometry regulates cancer cell tumorigenicity [75]. In comparison, our approach allows the mass production of shape-engineered multicellular structures without specialized bioprinting equipment, while also providing the opportunity to extract kinetic descriptors of multicellular aggregate formation.

Limitations of the present approach include the use of a cytoskeleton-modulating agent, which may introduce off-target or long-term effects on cell phenotype and function, despite the demonstrated partial reversibility under the conditions tested. In addition, the efficiency of generating sharply defined corners is constrained by the combined tolerances of 3D printing, agarose casting, and cell filling, which can lead to rounding of geometrical features. These technical limitations currently restrict the achievable sharpness and geometric fidelity of highly angular designs.

Complex aggregate architectures may offer distinct advantages in structural packing, alignment, and integration into modular tissue constructs or bioprinting grids. Engineered geometries could enhance space-filling efficiency, increase the rate of spheroid fusion due to enlarged contact interfaces (e.g., block–block versus sphere–sphere contact), and promote anisotropic tissue organization by imposing directional constraints during assembly. The proposed shape-engineered construct production strategy can be combined with emerging technologies for precise positioning of spheroids and organoids, such as digital acoustofluidic trapping techniques [76], thereby enabling controlled spatial arrangement, guided fusion, and hierarchical assembly of multicellular aggregates into complex tissue-like structures. Shape-engineered spheroids may also be used to pattern growth factor presentation or guide co-culture interactions in organoid systems. Investigating the feasibility and implications of this shape-engineering technology might open new pathways for the design of biofabricated tissues and functional *in vitro* models.

Here we demonstrated that modulation of the cytoskeleton is required to achieve complex spheroid shapes in addition to the use of microwells with the desired geometry, since otherwise surface tension drives the aggregates toward a final spherical shape. CytD at low concentrations showed the most pronounced effect among the treatments used. It binds to the barbed ends of actin filaments, preventing their elongation and promoting their disassembly, thereby disrupting the actin network’s ability to maintain tension [77,78]. Although Y-27632 has various effects on actin cytoskeleton organization and contractility via inhibition of Rho-associated kinase [19,79], here, as in previous studies, its effect on spheroid formation and visco-capillary velocity was modest [21,80], indicating that the cell contractility is not completely abolished and that some compensatory mechanisms are probably active. Blebbistatin produced similarly modest effects to those of Y-27632 [21,64,81]. The effects of Y-27632 and blebbistatin on spheroid fusion were analyzed in a previous study [68] and were shown to slow down, but not completely abolish fusion dynamics, in agreement with our data on aggregate formation. Interestingly, depending on the cell type, blebbistatin has been reported to either disrupt or increase enhance aggregate compaction, which was suggested to depend on the primary compaction mediator (cadherins or integrins) in a given cell type [82].

## Conclusions

We developed a microwell-based experimental approach combined with CFD modeling to quantify the visco-capillary velocity governing multicellular spheroid assembly. By modulating this parameter, we demonstrated a strategy for shape engineering of forming cell aggregates. Systematic comparison of circular, square, and triangular microwells identified circularity dynamics in square geometries as the most robust indicator of compaction kinetics, largely independent of cell seeding density. Integration of AFM and parallel-plate compression allowed us to relate aggregation kinetics to surface tension and effective viscosity, revealing that cell-type differences are predominantly driven by surface tension. The application of the CFD Volume of Fluid (VoF) method to model spheroid rounding provided simulations that closely matched experimental observations, offering a framework to simulate tissue dynamics. Pharmacological modulation of cytoskeletal contractility revealed that actin depolymerization significantly reduces surface tension and can be used to obtain cell aggregates with specific shapes reflecting the initial microwell geometry.

Overall, this integrated experimental–computational framework enables quantitative assessment of the physical parameters governing multicellular spheroid assembly. The approach is versatile and based on open-source CFD tools, making it well-suited for mechanobiological studies and drug screening applications targeting tissue morphogenesis. At the same time, several limitations should be acknowledged. The use of pharmacological cytoskeletal modulation may introduce long-term or off-target biological effects. The Newtonian fluid approximation represents a long-time effective description and does not capture short-time viscoelastic or active remodeling processes. Additionally, geometric fidelity of engineered shapes is constrained by printing resolution, agarose casting, cell filling, and intrinsic smoothing of sharp features during cell aggregation. Future work will focus on comprehensive 3D validation of engineered geometries, systematic analysis of long-term structural stability and remodeling dynamics, and evaluation of the functional applicability of shape-engineered constructs in tissue assembly, drug screening, and biofabrication contexts.

## Supporting information

STL files

## Acknowledgments

This work was supported by the Russian Science Foundation (grant No. 23-74-10113, https://rscf.ru/en/project/23-74-10113/) (microwells design, 3D printing, experiments on AFM and microtester, CFD modeling); and by the state assignment of the Ministry of Health of the Russian Federation (Theme No. NZAF-2024-0006) (cell cultures, imaging of spheroid formation); and performed using the equipment of the Shared Use Center “Center for Laser Technologies in Medicine” with the support of the Ministry of Science and Higher Education of the Russian Federation (Agreement No. 075-15-2025-669 dated August 5, 2025). The authors acknowledge Dr. Yanfang Geng and Dr. Mengliang Zhu from the CAS Key Laboratory for Biomedical Effects of Nanomaterials and Nanosafety, CAS Center for Excellence in Nanoscience, National Center for Nanoscience and Technology of China, for their assistance with data processing and manuscript proofreading.

