## Supplemental text for "Spheroid Assembly in Microwells of Defined Geometry for Quantitative Assessment of Aggregation Kinetics and Shape Engineering"


This supplementary documentation has been created to provide additional information to support the main text and contains:

Supplementary text “Comparison of SAMJ-IJ–based and manual segmentation of spheroids and assessment of the effective microwell geometry”

Supplementary text “Surface tension estimation by atomic force microscopy and parallel plate compression”

Supplementary text “Estimation of surface-to-volume ratio in different types of cell aggregates”

Supplementary figures S1 – S7

Supplementary movies S1 – S11

Supplementary .stl files 3D models of stamps (circular, square, and triangular)

**Supplementary text "Comparison of SAMJ-IJ–based and manual segmentation of spheroids and assessment of the effective microwell geometry"**

A subset of cell aggregates at selected time points was manually segmented to validate the SAMJ-IJ–based workflow (n = 20 aggregates per time point, 6 time points). The resulting time-dependent curves of projected area and circularity were highly similar between the two approaches (Fig. S2A,B).

For quantitative comparison, traced contours were converted into binary masks, and segmentation agreement between SAMJ-IJ and manual masks was assessed using the Dice similarity coefficient and Intersection over Union (IoU, Jaccard index). The Dice coefficient is defined as twice the intersection of two sets divided by the sum of their sizes. The IoU is calculated by dividing the area of overlap by the area of combined coverage. Across all spheroids and time points, the Dice coefficient was 0.94 ± 0.10 (mean ± SD), and the IoU was 0.92 ± 0.10, indicating strong agreement between the two segmentation methods (Fig. S2C,D).

Bland–Altman analysis was additionally performed for projected area and circularity. For projected area, SAMJ-IJ estimates were on average ~10% higher than manual measurements, likely due to improved detection of fine contour protrusions. Manual segmentation often implicitly smooths small boundary fluctuations due to hand-tracing limitations, which may lead to a slight underestimation of area. In addition, systematic dilation/erosion tendencies in one of the methods, potentially arising from thresholding differences at low-contrast boundaries, may contribute to the observed bias (Fig. S2E). For circularity, the mean bias was ~3%. A systematic trend was observed, with circularity values estimated by SAMJ-IJ being lower at high circularity and higher at low circularity, which may also reflect differences in perimeter tracing accuracy at early aggregation stages, when contours are more irregular (Fig. S2F).

We further assessed the sensitivity of the SAMJ-IJ–based workflow to the placement of the initial prompt by repeating segmentation on the same subset of spheroids (n = 20), using three independent sets of manually selected prompt positions within each aggregate in the first frame (Fig. S3A,B). The resulting projected area and circularity trajectories were virtually indistinguishable across prompt selections, demonstrating minimal dependence of the segmentation outcome on the exact prompt location.

To assess the effective microwell geometry resulting from the combined effects of stamp fabrication, agarose casting, and cell seeding, we performed shape analysis of the initial cell-filled microwell contours and compared them with the nominal CAD geometries using Dice similarity coefficients and Hausdorff distances (Fig. S3). The Hausdorff distance measures the largest minimal Euclidean distance between two contours, capturing the maximum local discrepancy between experimental and reference shapes. Specifically, for each point on one contour, the shortest distance to the other contour is determined, and the largest of these minimal distances is reported. Aligned contours from two independent cell-seeding experiments are shown in Fig. S3A, with the corresponding CAD geometries overlaid.

Dice scores were high for square and circular microwells (0.91 ± 0.03, n = 137, and 0.94 ± 0.02, n = 148, respectively), indicating strong global shape correspondence, but were lower for triangular (0.76 ± 0.03, n = 130) and star-shaped microwells (0.76 ± 0.07, n = 171) (Fig. S3B). Deviations were primarily localized at sharp corners, particularly in triangular and star-shaped microwells, where rounding consistent with printing and casting limits was observed. These local discrepancies were reflected in the Hausdorff distance values, which were low for circular (38 ± 8 μm) and square (45 ± 10 μm) geometries, and higher for triangular (118 ± 11 μm) and star-shaped (107 ± 26 μm) geometries, as expected due to their sharper features.

**Supplementary text "Surface tension estimation by atomic force microscopy and parallel plate compression"**

The surface tension of spheroids was estimated using a combined approach established in our previous work (Kosheleva et al., 2023). In this method, atomic force microscopy (AFM) provides measurements of the local elasticity of the spheroid's outer cell layer, while parallel plate compression reflects the global mechanical response of the entire spheroid. By comparing the two, surface tension can be isolated, as it represents the fraction of total compressive resistance arising from interfacial forces rather than bulk deformation.

AFM measurements were performed using a Bioscope Resolve system (Bruker, Santa Barbara, CA) mounted on an Axio Observer inverted optical microscope (Carl Zeiss, Germany). The sample temperature was maintained at 37 °C throughout the experiments using a heated stage. Spheroids were transferred in growth medium onto cell culture Petri dishes and were incubated for 1–2 hours to promote adhesion. Due to the spheroidal geometry, AFM measurements were taken from the central top region of each spheroid, where the surface was relatively flat. For nanomechanical mapping, PeakForce QNM-Live Cell probes (PFQNM-LC-A-CAL, Bruker AFM Probes, Camarillo, CA, USA) were used. These probes feature short paddle-shaped cantilevers with pre-calibrated spring constants in the range of 0.09–0.11 N/m and 17 μm-long tips with a nominal radius of 70 nm. Fast Force Volume mode was employed to acquire nanomechanical maps, with a spatial resolution of 40x40 points over an area of 60 × 60 $\mu m^2$. Force curves were acquired at a vertical piezo speed of 180 μm/s (30 Hz ramp rate, 3 μm ramp size) with a force setpoint of 0.5–1 nN, resulting in indentation depths of

~500 nm. The individual force curves from the volume maps were processed with the Python scripts, as described previously (Efremov et al., 2019), using the Hertz's model:

$$F(\delta) = \frac{4\sqrt{R_{tip}}}{3(1-\nu^2)} E_{ind} \delta^{\frac{3}{2}}; \quad \text{(S1)}$$

where $F$ is the force acting on the cantilever; $\delta$ is the indentation depth; $\nu$ is the Poisson's ratio of the sample (assumed to be equal to 0.5); $R_{tip}$ is the tip radius; and $E_{ind}$ is the effective Young's indentation modulus. For each spheroid, the $E_{ind}$ values were averaged across the full force map to obtain the mean modulus. Measurements were performed on at least 10 spheroids per cell type. The model provided an excellent fit to the experimental data, with an adjusted $R^2 > 0.97$ (Fig. S4A).

Parallel-plate compression was performed with a micro-scale mechanical testing system (Microsquisher, CellScale, Canada) using the SquisherJoy software. Spheroids were placed in a PBS-filled bath and compressed between a fixed rigid substrate and a movable cantilever beam equipped with a flat platform. Each spheroid was compressed up to 30% of its initial height at a displacement speed of 10 μm/s, while simultaneously recording the applied force, the upper plate position, and real-time spheroid deformation using a side-view camera. At least 10 spheroids per cell type were analyzed. The mechanical model for spheroid compression from (Kosheleva et al., 2023) was applied to estimate the bulk Young's modulus ($E_{bulk}$). In this model, a multiplicative explicit correction factor $f(\delta)$ expressed as a function of the normalized indentation depth ($\delta/R_{sph}$, where $R_{sph}$ is the radius of the spheroid) was introduced:

$$F = \frac{4}{3} f(\delta) \frac{E_{bulk}}{1-\nu^2} \delta^{\frac{3}{2}} \sqrt{R_{sph}}; \quad \text{(S2)}$$

$$f(\delta/R_{sph}) = 1 + d_1\left(\frac{\delta}{R_{sph}}\right)^{\frac{1}{2}} + d_2\left(\frac{\delta}{R_{sph}}\right) + d_3\left(\frac{\delta}{R_{sph}}\right)^{\frac{3}{2}} + d_4\left(\frac{\delta}{R_{sph}}\right)^{2} + d_5\left(\frac{\delta}{R_{sph}}\right)^{\frac{5}{2}}. \quad \text{(S3)}$$

The radius of the spheroid $R_{sph}$ was assessed with the side-view camera during the measurements. The coefficients of the correction function $f(\delta)$ were obtained by fitting finite element method (FEM) simulation curves, as described previously (Kosheleva et al., 2023). To characterize the relaxation time of the cell spheroids in compression experiments, the Lee–Radok model was used to process both the approach and force relaxation phases as follows:

$$F = \frac{4}{3}\frac{\sqrt{R}}{1-\nu^2} f(\delta) \int_0^t E(t-\xi)\frac{\partial \delta^{\frac{3}{2}}}{\partial \xi} d\xi; \quad \text{(S4)}$$

where $E(t)$ is the Young's relaxation modulus for the selected viscoelastic model, and $\xi$ is the dummy time variable required for the integration. The numerical processing was performed using the previously developed MATLAB scripts [82]. The standard linear solid (SLS) viscoelastic model was applied as the relaxation modulus function, yielding excellent fit quality with an adjusted $R^2 > 0.97$ and allowing estimation of the characteristic relaxation time.

By combining measurements of surface elasticity ($E_{ind}$) and bulk elasticity ($E_{bulk}$), the effective surface tension was calculated based on the model proposed by Ding et al. (Ding et al., 2015). This model provides an explicit expression that links the loading force to indentation depth:

$$F = \frac{4}{3}\frac{E_{bulk}}{1-\nu^2}\delta^{\frac{3}{2}}\sqrt{R_{tip}}\left(1 + 1.889\left(\frac{s}{\delta}\right)^{0.437}\left(\frac{s}{R_{tip}}\right)^{0.5}\right); \quad \text{(S5)}$$

$$s = \frac{\sigma}{E_{bulk}}(1-\nu^2); \quad \text{(S6)}$$

where the parameter $s$ links the surface tension σ and the bulk Young's modulus $E_{bulk}$. In parallel-plate compression, where the entire spheroid is deformed, the influence of surface energy is negligible, and $E_{bulk}$ is directly obtained. However, in AFM experiments with a small indenter radius (≈100 nm) and shallow indentation depth (≈500 nm), the contribution of surface tension becomes significant. Under these conditions, the indentation modulus relates to the bulk modulus as:

$$E_{ind} = E_{bulk}\left(1 + 1.889\left(\frac{s}{\delta}\right)^{0.437}\left(\frac{s}{R_{tip}}\right)^{0.5}\right); \qquad \text{(S7)}$$

By inserting the mean values for the indenter radius, indentation depth, and the experimentally determined bulk and surface elastic moduli for the two spheroid types, we estimated $s$ and thereby surface tension σ.

Representative AFM force maps and force curves are shown in Fig. S4A. The effective indentation modulus $E_{ind}$ was significantly different between cell types: 4.4 ± 1.2 kPa (n=10) for ARPE-19 spheroids versus 12.4 ± 2.1 kPa (n=16) for HDF spheroids ($p<0.001$, Mann–Whitney test). Parallel-plate compression yielded consistently lower bulk elastic moduli ($E_{bulk}$): 1.9 ± 0.9 kPa (n=23) for ARPE-19 and 2.7 ± 1.1 kPa (n=20) for HDF spheroids ($p<0.05$) (Fig. S4B). By combining these values (Fig. S4C) and applying Eqs. S4–S6, the estimated surface tension was 0.27 ± 0.15 N/m for ARPE-19 spheroids and 1.15 ± 0.5 N/m for HDF spheroids. The parameters are summarized in Table S1.

**Supplementary text "Estimation of surface-to-volume ratio in different types of cell aggregates"**

The spherical shape exhibits the minimal surface-to-volume ratio (S/V), given by:

$$S = 4\,\pi R^2\ ; \tag{S8}$$

$$V = \frac{4}{3}\,\pi R^3; \tag{S9}$$

$$\frac{S}{V} = \frac{3}{R}\ . \tag{S10}$$

For the brick-like aggregates, assuming a cube-like rectangular prism with base side length *a* and thickness *h*:

$$S_b = 2(a^2 + 2ah)\ ; \tag{S11}$$

$$V_b = a^2 h\ ; \tag{S12}$$

$$\frac{S_b}{V_b} = \frac{2(a+2h)}{ah} = 2\left(\frac{1}{h} + \frac{2}{a}\right). \tag{S13}$$

For a typical spherical spheroid with a diameter of 150 μm, and assuming the same volume for a brick-like aggregate with a thickness of 50 μm (approximate value taken from the confocal Z-stacks), the latter exhibits an approximately 50% higher surface-to-volume ratio (0.061 vs 0.04 $\mu m^{-1}$).

For a star-shaped aggregate, which can be approximated as a star-shaped prism of thickness *h*, the general expression for the surface-to-volume ratio is:

$$S = 2A_{base} + P_{base}h\ ; \tag{S14}$$

$$V = A_{base}h\ ; \tag{S15}$$

$$\frac{S}{V} = \frac{2}{h} + \frac{P_{base}}{A_{base}}; \tag{S16}$$

where $A_{base}$ is the area of the base and $P_{base}$ is its perimeter. For a 5-point star, the perimeter is typically 1.5–2× that of a circle or square with the same area. Considering conservative case of $P_{star} = 1.6P_{square}$, the same thickness (50 µm) and the same volume, we obtain around a 2× increase relative to the surface-to-volume ratio of spherical aggregates (0.074 vs 0.04 $\mu m^{-1}$).

### Supplementary Tables

**Table S1.** Parameters used in spheroid mechanical analysis, their designations and values.

| Parameter | Designation | Value | |
|---|---|---|---|
| | | ARPE | HDF |
| Poisson's ratio of spheroid (from literature) | $\nu$ | 0.5 | |
| AFM tip radius (from manufacturer) | $R_{tip}$ | 70 nm | |
| Indentation depth in AFM experiments | $\delta$ | 500-1000 nm | |
| Effective Young's indentation modulus | $E_{ind}$ | 4.4 ± 1.2 kPa | 12.4 ± 2.1 kPa |
| Radius of spheroid from parallel plate compression experiments | $R_{sph}$ | 180 ± 30 μm | 160 ± 30 μm |
| Bulk spheroid Young's modulus from parallel plate compression experiments | $E_{bulk}$ | 1.9 ± 0.9 kPa | 2.7 ± 1.1 kPa |
| Surface tension | σ | 0.27 ± 0.12 N/m | 1.15 ± 0.5 N/m |
| Relaxation time | $\tau_{rel}$ | 2.7 ± 0.8 s | 4.3 ± 1.1 s |
| Visco-capillary velocity | $v_p$ | 22 ± 4 μm/s | 3.9 ± 0.4 μm/s |
| Effective viscosity | η | $(5.0\pm0.1)\cdot10^4$ Pa·s | $(7.3\pm0.1)\cdot10^4$ Pa·s |

**Table S2.** Shape descriptors of ARPE-19 and HDF spheroids formed in square microwells with different treatments affecting surface tension (cytochalasin D (cytD), blebbistatin (blebb), and Y-27632). Shape descriptors: circularity, roundness, solidity, perimeter-to-normalized-area ratio, maximum curvature ($\kappa_{max}$), curvature variance (Var(κ)).

| Spheroid type | Treatment | Circularity | Roundness | Solidity | Maximum curvature ($\kappa_{max}$) | Curvature variance (Var(κ)) | Perimeter-to normalized-area ratio |
|---|---|---|---|---|---|---|---|
| ARPE-19 | Control (n=116) | 0.894 ± 0.007 | 0.48 ± 0.01 | 0.987 ± 0.002 | 2.10 ± 0.10 | 0.24 ± 0.04 | 2.115 ± 0.008 |
| | cytD, 0.5 μM (n=160) | 0.88 ± 0.01 | 0.47 ± 0.02 | 0.986 ± 0.003 | 2.4 ± 0.2 | 0.38 ± 0.08 | 2.14 ± 0.02 |
| | blebbistatin, 10 μM (n=160) | 0.890 ± 0.008 | 0.47 ± 0.01 | 0.986 ± 0.002 | 2.1 ± 0.1 | 0.27 ± 0.04 | 2.12 ± 0.01 |
| | Y-27632, 50 μM (n=120) | 0.896 ± 0.007 | 0.47 ± 0.01 | 0.987 ± 0.002 | 2.0 ± 0.1 | 0.23 ± 0.03 | 2.113 ± 0.009 |
| HDF | control (n=135) | 0.891 ± 0.007 | 0.46 ± 0.02 | 0.986 ± 0.002 | 2.1 ± 0.1 | 0.25 ± 0.04 | 2.119 ± 0.009 |
| | cytD, 1 μM (n=100) | 0.83 ± 0.02 | 0.46 ± 0.02 | 0.977 ± 0.008 | 3.5 ± 0.5 | 1.0 ± 0.2 | 2.20 ± 0.03 |
| | blebbistatin, 10 μM (n=146) | 0.893 ± 0.007 | 0.46 ± 0.02 | 0.987 ± 0.002 | 2.03 ± 0.10 | 0.23 ± 0.03 | 2.116 ± 0.008 |
| | Y-27632, 50 μM (n=149) | 0.899 ± 0.007 | 0.48 ± 0.01 | 0.988 ± 0.002 | 2.0 ± 0.1 | 0.21 ± 0.03 | 2.110 ± 0.008 |

**Table S3.** Shape descriptors of ARPE-19 and HDF spheroids formed in various microwell geometries (circular, square, triangular, and star-shaped) with cytD treatment (except for circular microwells, which were incubated with DMSO). Shape descriptors: circularity, roundness, solidity, perimeter-to-normalized-area ratio, maximum curvature ($\kappa_{max}$), curvature variance (Var(κ)).

| Spheroid type | Type of microwells | Circularity | Roundness | Solidity | Maximum curvature ($\kappa_{max}$) | Curvature variance (Var(κ)) | Perimeter-to-normalized-area ratio |
|---|---|---|---|---|---|---|---|
| ARPE-19 | circular (n=116) | 0.896 ± 0.007 | 0.47 ± 0.01 | 0.986 ± 0.002 | 2.00 ± 0.10 | 0.22 ± 0.03 | 2.113 ± 0.008 |
| | square (n=159) | 0.87 ± 0.02 | 0.47 ± 0.02 | 0.985 ± 0.003 | 2.4 ± 0.2 | 0.40 ± 0.10 | 2.14 ± 0.02 |
| | triangular (n=61) | 0.77 ± 0.04 | 0.44 ± 0.02 | 0.97 ± 0.02 | 3.8 ± 0.9 | 1.0 ± 0.4 | 2.28 ± 0.06 |
| | star (n=169) | 0.82 ± 0.03 | 0.46 ± 0.02 | 0.96 ± 0.01 | 3.4 ± 0.5 | 0.9 ± 0.3 | 2.21 ± 0.04 |
| HDF | circular (n=139) | 0.88 ± 0.01 | 0.44 ± 0.03 | 0.986 ± 0.002 | 2.1 ± 0.1 | 0.26 ± 0.04 | 2.13 ± 0.01 |
| | square (n=82) | 0.81 ± 0.03 | 0.45 ± 0.02 | 0.97 ± 0.02 | 3.8 ± 0.9 | 1.1 ± 0.5 | 2.23 ± 0.04 |
| | triangle (n=116) | 0.74 ± 0.03 | 0.43 ± 0.02 | 0.97 ± 0.01 | 4.1 ± 0.5 | 1.3 ± 0.2 | 2.33 ± 0.04 |
| | star (n=108) | 0.72 ± 0.04 | 0.46 ± 0.02 | 0.90 ± 0.03 | 5.1 ± 0.9 | 1.9 ± 0.6 | 2.36 ± 0.07 |

### Supplementary Figures

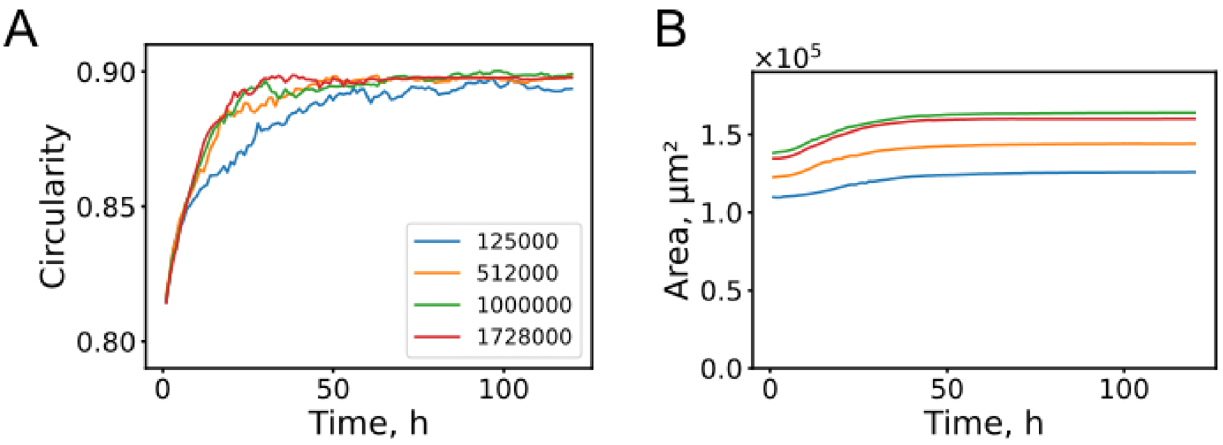


**Figure S1. Mesh sensitivity analysis in OpenFOAM for extracted morphological parameters.** A cuboidal aggregate model (400 × 400 × 400 μm) was simulated within a computational domain of 1200 × 1200 × 1200 μm. (A) Circularity. (B) Projected surface area. The number of mesh elements used in each simulation is indicated in the legend.

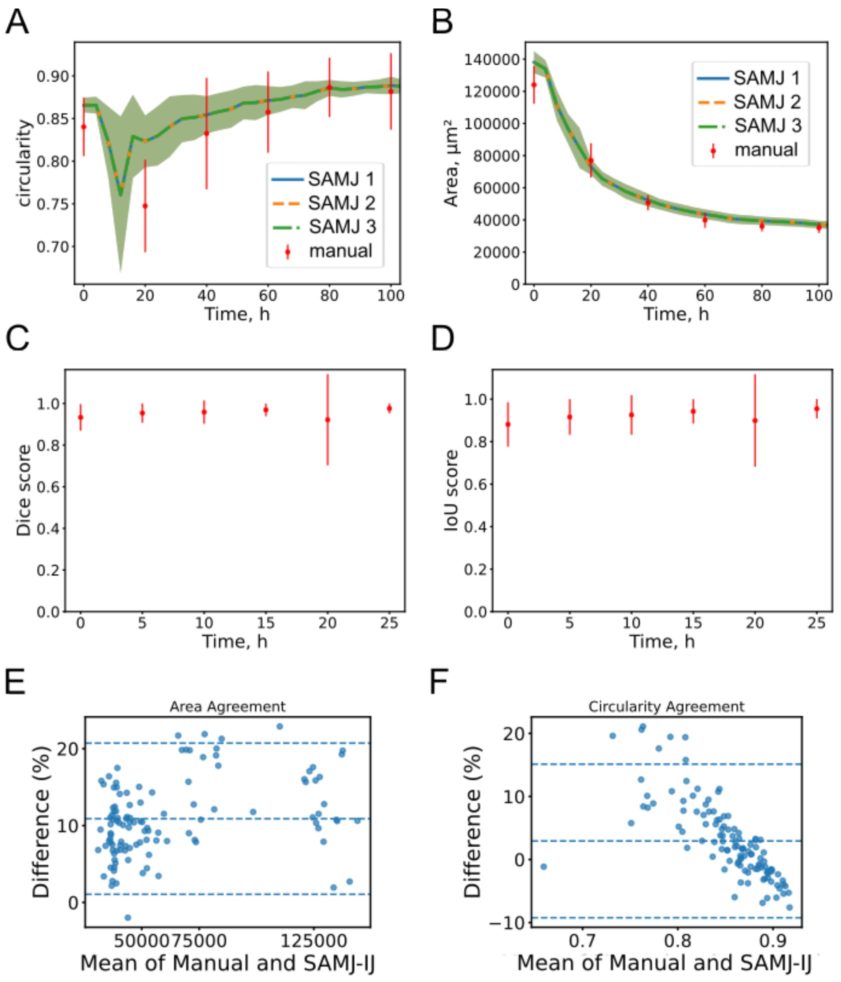


**Figure S2. Comparison of spheroid segmentation using the SAMJ-IJ plugin and manual segmentation.** (A, B) Time-dependent circularity and projected surface area for 20 spheroids analyzed across five time points. Scatter plots with standard deviation represent manual segmentation results. Continuous curves correspond to SAMJ-IJ–based segmentation performed using three independent sets of manually selected prompt positions (SAMJ 1, SAMJ 2, SAMJ 3); shaded regions indicate standard deviation. (C, D) Dice similarity coefficient and Intersection over Union (IoU, Jaccard index) quantifying agreement between SAMJ-IJ and manually generated masks. (E, F) Bland–Altman plots comparing projected area and circularity measurements obtained by SAMJ-IJ and manual segmentation.

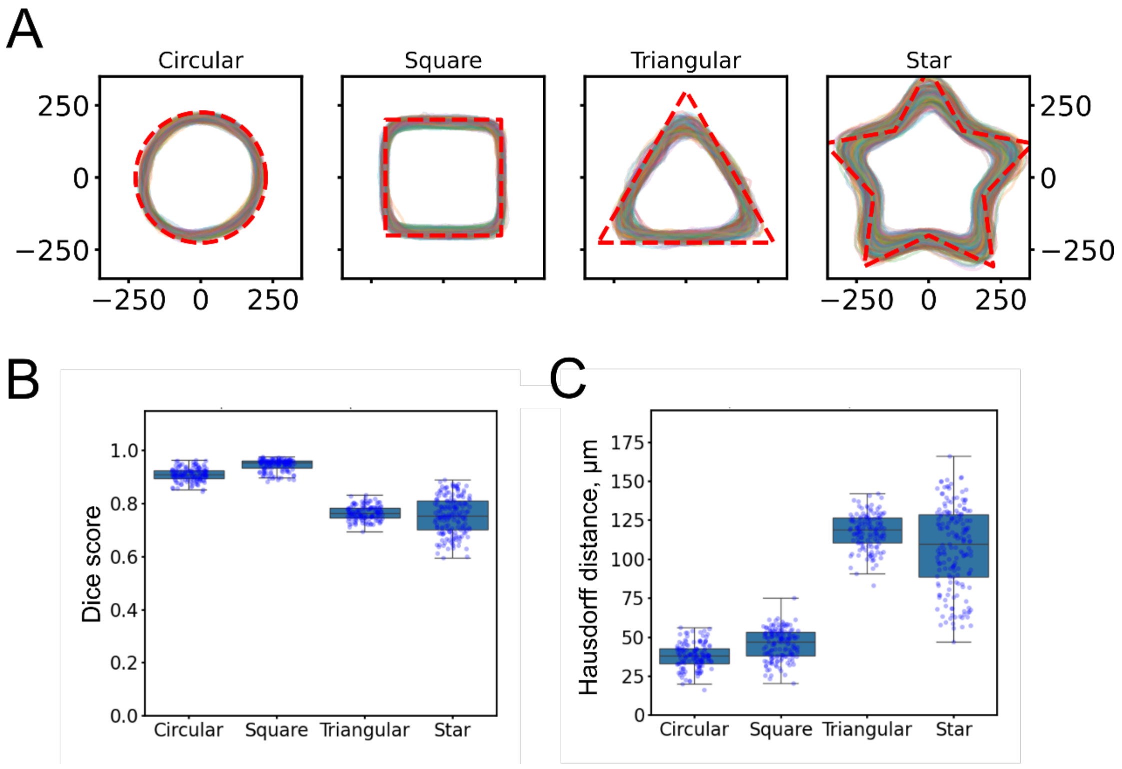


**Figure S3. Assessment of effective microwell geometry based on initial cell-filled microwell contours.** (A) Representative aligned contours of initial cell-filled microwells obtained using SAMJ-IJ–based segmentation for different geometries. Nominal CAD geometries used for stamp fabrication are overlaid as dotted lines. (B) Dice similarity coefficients quantifying global shape agreement between experimental contours and corresponding CAD geometries. (C) Hausdorff distances quantifying the maximum local geometric deviation between experimental contours and corresponding CAD geometries.

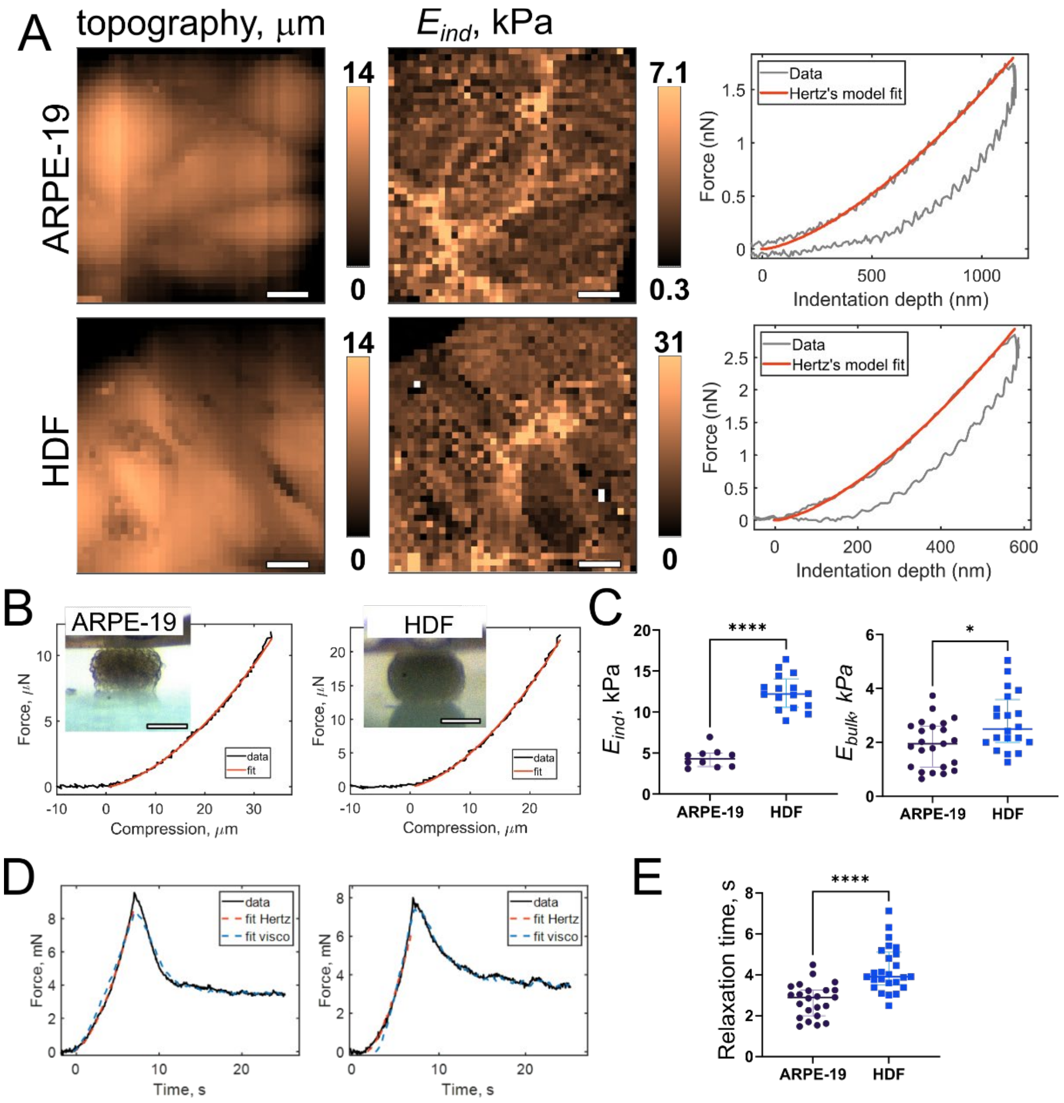


**Figure S4. Mechanical analysis of ARPE-19 and HDF spheroids. (A)** Representative AFM force maps showing topography and effective Young's indentation modulus ($E_{ind}$); scale bar: 10 µm. Example force–distance curves with corresponding Hertz model fits are shown. (B) Force–compression curves obtained from parallel-plate compression experiments with the fitted Hertzian mechanical model. (C) Quantitative comparison of Young's indentation modulus ($E_{ind}$) measured by AFM and bulk modulus ($E_{bulk}$) obtained from compression tests. Each dot represents a single spheroid; horizontal lines indicate median values and interquartile ranges. (D) Compression data plotted as force versus time during stress relaxation, with standard linear solid (SLS) model fits. (E) Relaxation times of ARPE-19 and HDF spheroids derived from compression experiments.

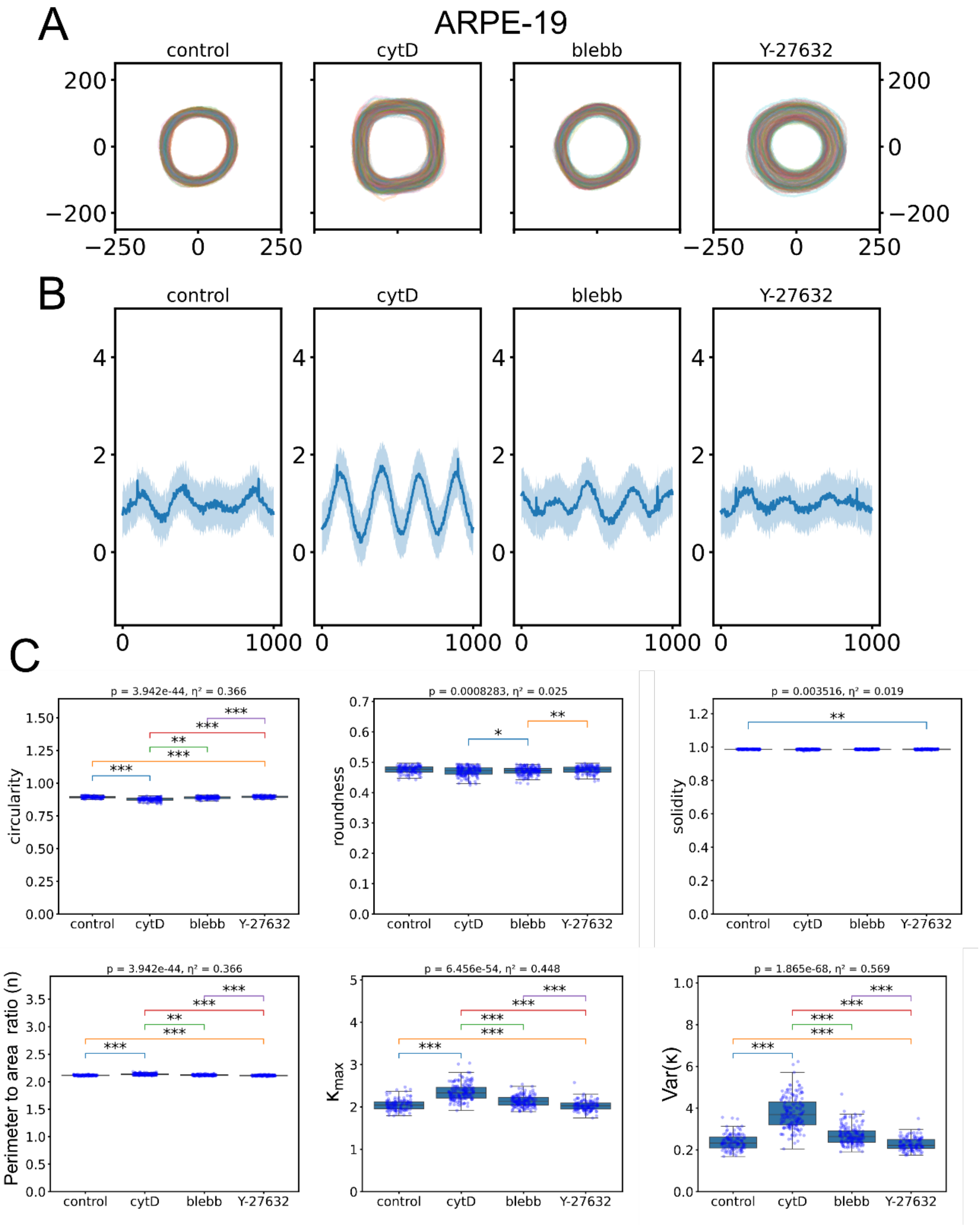


**Figure S5. Shape descriptors of ARPE-19 spheroids after treatments affecting surface tension (cytochalasin D (cytD), blebbistatin (blebb), and Y-27632). (A)** Aligned contours **(B)** Local boundary curvature. **(C)** Shape descriptors: circularity, roundness, solidity, perimeter-to-normalized-area ratio, maximum curvature ($\kappa_{max}$), curvature variance (Var(κ)) (*p < 0.05, **p < 0.01, ***p < 0.001).

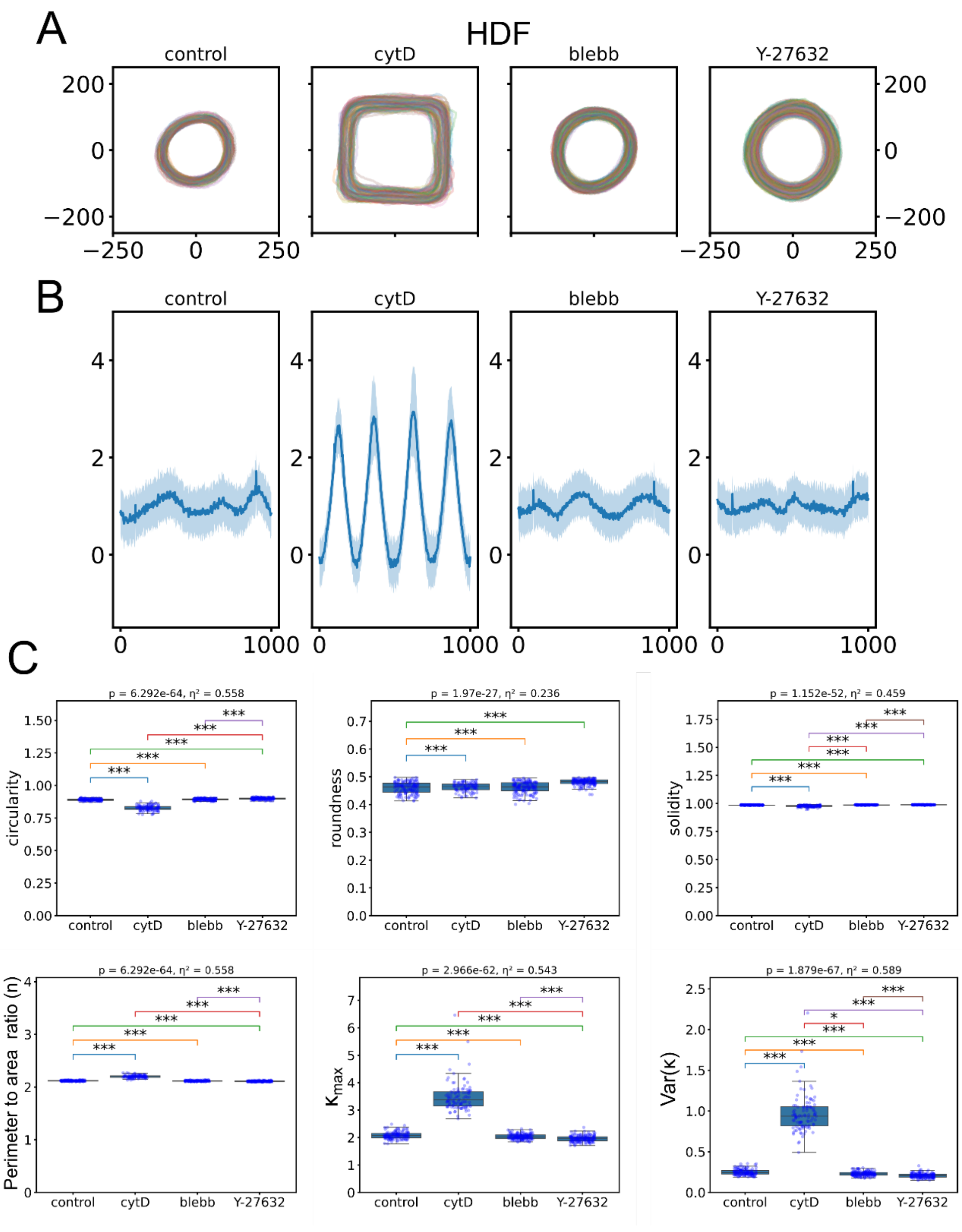


**Figure S6. Shape descriptors of HDF spheroids after treatments affecting surface tension (cytochalasin D (cytD), blebbistatin (blebb), and Y-27632). (A)** Aligned contours **(B)** Local boundary curvature. **(C)** Shape descriptors: circularity, roundness, solidity, perimeter-to-normalized-area ratio, maximum curvature ($\kappa_{max}$), curvature variance (Var(κ)) (*p < 0.05, **p < 0.01, ***p < 0.001).

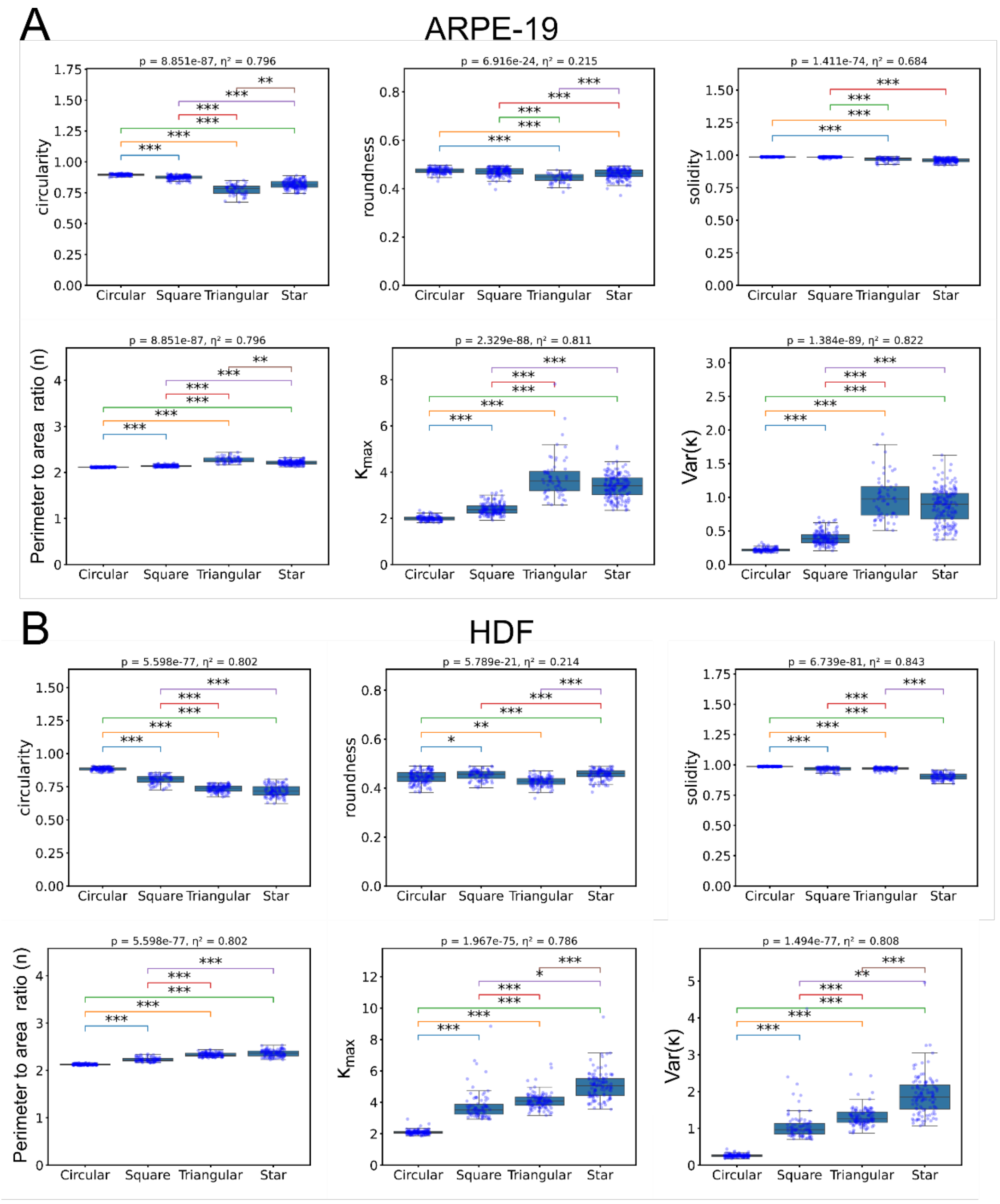


**Figure S7. Shape descriptors of shape-engineered ARPE-19 and HDF spheroids (circular, brick-like, prismatic, and star-shaped).** (A) ARPE-19 spheroids, shape descriptors: circularity, roundness, solidity, perimeter-to-normalized-area ratio, maximum curvature ($\kappa_{max}$), curvature variance (Var(κ)) (*$p < 0.05$, **$p < 0.01$, ***$p < 0.001$). (B) HDF spheroids, shape descriptors: circularity, roundness, solidity, perimeter-to-normalized-area ratio, maximum curvature ($\kappa_{max}$), curvature variance (Var(κ)) (*$p < 0.05$, **$p < 0.01$, ***$p < 0.001$).

**Supplementary Movies**

**Movie S1.** Example of spheroid formation dynamics, shown as a stitched time-lapse from a scan over a single well of a 24-well plate, acquired using the CellInsight CX7 High-Content Screening (HCS) Platform (Thermo Fisher Scientific). Case: ARPE-19 cell aggregation in square microwells. Time step: 4 h.

**Movie S2.** Formation of spheroids from ARPE-19 and HDF cells in microwells of different cross-sectional geometries (circular, square, triangular). A single representative spheroid was selected for each case. Time step: 4 h.

**Movie S3.** Example of SAM-based segmentation of ARPE-19 cell aggregates in square microwells. Left: original movie; right: ROIs from segmentation output (yellow outlines). Time step: 4 h.

**Movie S4.** CFD simulation results using the VoF method in OpenFOAM for an initial cylindrical geometry (circular cross-section). Shown are an angled view and three projections (top to bottom: ZX, ZY, XY).

**Movie S5.** CFD simulation results using the VoF method in OpenFOAM for an initial rectangular cuboid geometry (square cross-section). Shown are an angled view and three projections (top to bottom: ZX, ZY, XY).

**Movie S6.** CFD simulation results using the VoF method in OpenFOAM for an initial triangular prism geometry (triangular cross-section). Shown are an angled view and three projections (top to bottom: ZX, ZY, XY).

**Movie S7.** Direct comparison of experimental spheroid formation data and OpenFOAM simulation results. Case: ARPE-19 cell aggregation in square microwells. OpenFOAM simulation (XY projection) with the same geometrical parameters and a visco-capillary velocity of 10 μm/s. Time step: 4 h.

**Movie S8.** Modulation of ARPE-19 spheroid formation by treatments affecting surface tension. Left to right: control, cytochalasin D (cytD), blebbistatin (blebb), and Y-27632.

Time-lapse observation of ARPE-19 spheroid formation in square microwells. Time step: 4 h; scale bar: 200 μm.

**Movie S9.** Modulation of HDF spheroid formation by treatments affecting surface tension. Left to right: control, cytochalasin D (cytD), blebbistatin (blebb), and Y-27632. Time-lapse observation of HDF spheroid formation in square microwells. Time step: 1.5 h; scale bar: 200 μm.

**Movie S10.** Formation of brick-shaped spheroids from ARPE-19 cells in square microwells under cytochalasin D (cytD) treatment. Control spheroids formed under standard conditions are shown for comparison. Time step: 4 h; scale bar: 100 μm.

**Movie S11.** Formation of star-shaped spheroids from ARPE-19 cells in star-shaped microwells under cytochalasin D (cytD) treatment. Control spheroids formed under standard conditions are shown for comparison. Time step: 4 h; scale bar: 100 μm.
